# Multi-omic Environmental Monitoring Reveals When Molecular Signals Reflect Living Biology

**DOI:** 10.64898/2026.09.04.749360

**Authors:** Yin Cheong Aden Ip, Gledis Guri, Pedro FP Brandão-Dias, Elizabeth Andruszkiewicz Allan, Ryan P. Kelly

## Abstract

Environmental molecular monitoring has transformed biodiversity detection, yet most methods capture only species presence through amplified DNA fragments, missing the biological information encoded in native nucleic acids. Here we show that a single shotgun Oxford Nanopore sequencing library from water can be read through several independent biological lenses, and that each class of molecule (environmental RNA, CpG methylation, and bulk DNA) differs in its environmental persistence, which sets the window over which signal can be trusted. Sampling Pacific salmon spawning runs, we simultaneously recovered environmental RNA indicating fresh biological input, CpG methylation profiles reflecting age demographics, and pathogen signatures reflecting health. Environmental RNA tracked spawning phenology, qPCR corroborated the timing of fresh biological input over the weeks sampled, and methylation-based age inference held only while fresh DNA dominated, a condition we term the’Freshness Gate.’ Because environmental water is not a tissue sample, we detect functional transcripts as evidence of biological input but do not infer differential gene expression, and we read methylation state but do not infer biological age outside the Freshness Gate. Variance partitioning is consistent with biological freshness, rather than technical factors such as fragment length or sequencing depth, structuring environmental methylation, though these comparisons did not reach significance at nine time points. Comparable dynamics were observed in Chinook salmon, with timing offsets following their distinct spawning season. By defining when environmental molecules faithfully encode living biology and when degradation decouples signal from current state, our framework sets practical and theoretical boundaries for multi-omic environmental monitoring.

## 1. Introduction

Environmental molecular monitoring has revolutionized biodiversity assessment by enabling non-invasive species detection from trace DNA in water, soil, and air (Ficetola et al., 2008; Thomsen & Willerslev, 2015; Taberlet et al., 2018). These approaches predominantly rely on PCR amplification of short barcode regions (i.e., metabarcoding), which efficiently answers the fundamental question of which species are present in an ecosystem (Deiner et al., 2017; Goldberg et al., 2016). However, amplification-based methods also inherently select for specific gene targets, erase epigenetic modifications, and exclude non-target organisms entirely. This creates a fundamental limitation where they capture a constrained slice of taxonomic composition but miss the rich biological information encoded in native nucleic acid states (Barnes & Turner, 2016; Cristescu & Hebert, 2018), from physiological activity to pathogen communities that coexist in the same environmental sample (Figure 1).

**Figure 1.**
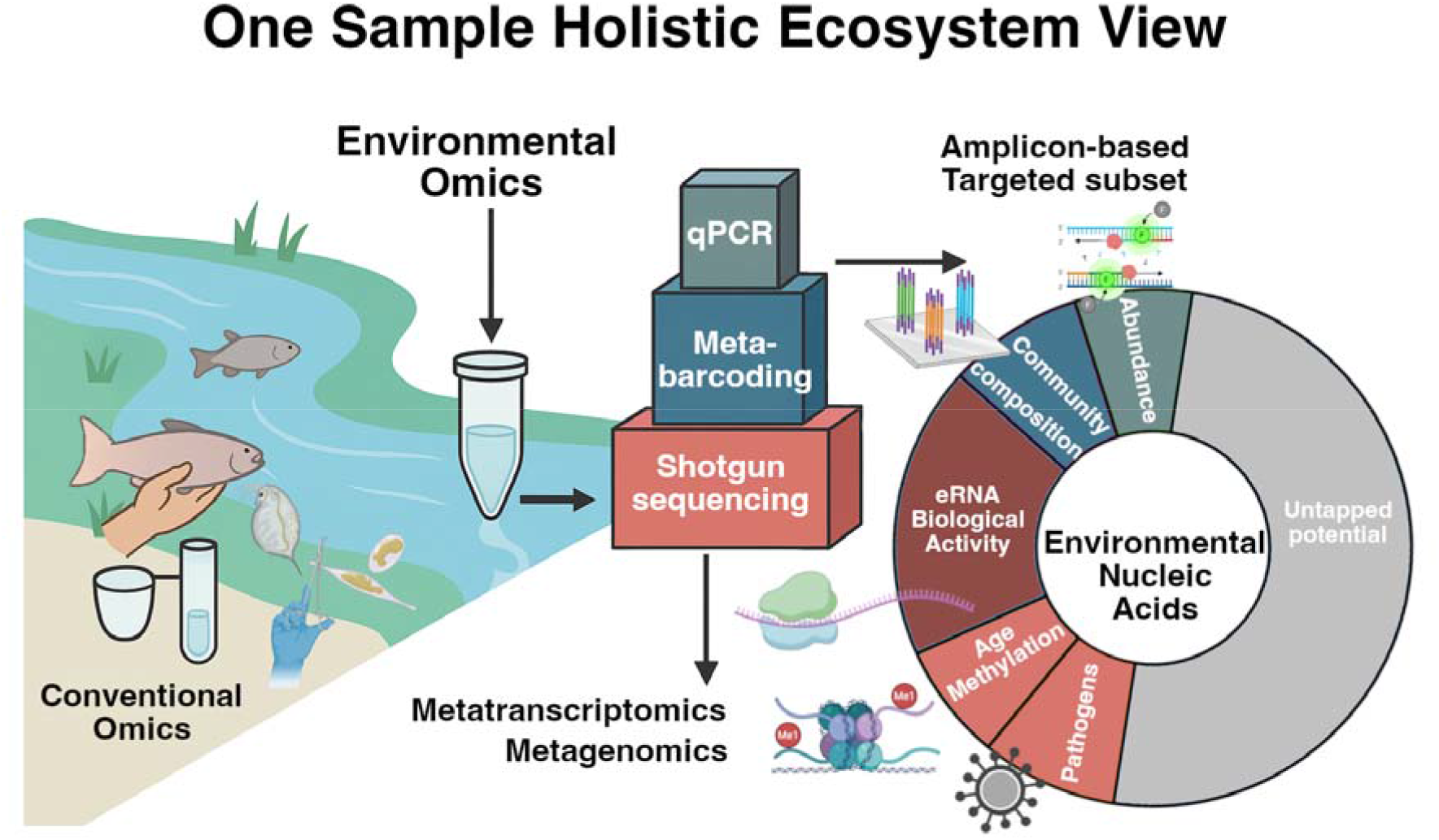
One Sample, Holistic Ecosystem View. Environmental omics approaches extract multiple biological signals from a single environmental sample, integrating qPCR, metabarcoding, RNA and DNA shotgun sequencing to generate complementary insights. Amplicon-based methods target known taxa to assess community composition and relative abundance, while shotgun sequencing enables metatranscriptomic and metagenomic analyses that reveal environmental RNA biological activity, age-related methylation patterns, and pathogen profiles. Together, these layers capture the full potential of native nucleic acids (DNA and RNA) to link biodiversity, physiological state, population demographics, and ecosystem health within a unified environmental monitoring framework.

Native environmental nucleic acids carry multiple layers of biological information beyond mere presence. Messenger RNA transcripts report real-time physiological activity and stress responses (Cristescu, 2019; Hechler et al., 2025). DNA methylation patterns encode developmental stages, age and demographic structure (Horvath, 2013; Hannum et al., 2013; Hirayama et al., 2024; Ruiz et al., 2025). Pathogen nucleic acids reveal ecosystem health dynamics (Bass et al., 2015; Ip et al., 2024; Ip et al., 2025b). Yet despite the theoretical richness of these signals, practical recovery of multi-omic information from complex environmental samples remains largely unexplored in studies focused on macroorganisms (Balard et al., 2024; Minamoto et al., 2022). Most animal-focused environmental RNA (eRNA) and epigenetic studies have been conducted in controlled mesocosm settings, largely due to the unanswered question: When do environmental molecules reliably reflect the current biological state, and when has degradation rendered them misleading?

In microbial and planktonic studies, environmental meta-transcriptomics and meta-epigenomics have successfully revealed community-level functional states, diel cycles, and nutrient response patterns in field-collected environmental samples (Blaskowski et al., 2025; Lambert et al., 2022; Ottesen et al., 2014). These successes build on the fundamental characteristics of unicellular systems: abundant biological material, rapid cellular turnover, and limited spatial structure. Marine phytoplankton communities, for instance, can be comprehensively sampled from water volumes that simultaneously capture millions of intact cells (Marchetti et al., 2012), enabling robust inference of gene expression and epigenetic regulation at the ecosystem scale (e.g., Alexander et al., 2015; Bertrand et al., 2015).

Extending this environmental omics framework to multicellular animals introduces several different challenges. First, the target organisms themselves are sparse and patchily distributed. Additionally, environmental DNA (eDNA) is derived from heterogeneous sources, such as sloughed epithelial cells, gametes, excreta, decaying tissue; each with distinct molecular properties and degradation kinetics. Environmental RNA (eRNA) degrades rapidly outside cells, typically within hours to days in aquatic environments (Yates et al., 2021; Cristescu, 2019). Most critically, environmental samples capture a mixture of fresh biological input (both eDNA and eRNA) and accumulated degraded material, with their relative contributions changing continuously as organisms arrive, spawn, and senesce. These complexities mean that the temporal window for reliable environmental inference may be narrow, system-dependent, and time-dependent.

Recent sequencing advances by Oxford Nanopore Technologies (ONT) have created new opportunities for environmental multi-omics approaches (Ip et al., 2026; Simpson et al., 2017; Payne et al., 2021). Unlike short-read platforms that fragment DNA and strip molecular modifications during library preparation (Hirayama et al., 2024), nanopore sequencing directly interrogates native nucleic acid molecules—long reads spanning kilobases enable complete mitogenome assembly and haplotype phasing (Matthews et al., 2026; Mizuno et al., 2026), real-time modified basecalling detects DNA methylation states without bisulfite treatment (Liu et al., 2019; Ni et al., 2019; Ruiz et al., 2025), and adaptive sampling allows real-time enrichment of target taxa while retaining unbiased background reads for community context (Payne et al., 2021).

These capabilities allow a single environmental sample to simultaneously yield eRNA for functional transcript detection, DNA methylation patterns for age demographics, and shotgun metagenomics for pathogen detection, all from the same library preparation and sequencing run. Until now, however, environmental RNA studies have relied on targeted qPCR of single transcripts or short-read RNA-seq that loses long-range information (e.g., Hechler et al., 2025; Miyata et al., 2021; Tsuri et al., 2021), while methylation profiling has been limited to bisulfite approaches that destroy native DNA and require large amounts of input material (Hirayama et al., 2024; Balard et al., 2024; Zhao et al., 2023). The convergence of ONT with environmental sampling creates a new opportunity to test what is realistically recoverable from environmental systems.

Pacific salmon provide an attractive model system for testing the applicability of environmental omics from macroorganism populations (Duda et al., 2021; Quinn, 2018). Anadromous salmon exhibit predictable seasonal migrations and synchronous spawning pulses, creating temporal structure in biological inputs (Groot & Margolis, 1991). Within a single watershed, we observe sequential phases: upstream migration, concentrated spawning aggregations depositing massive amounts of gametes and somatic tissue, and post-spawning senescence producing carcass material (Biela et al., 2022). These life-history dynamics generate natural controlled experiments where biological inputs transition from predominantly fresh (i.e., actively spawning adults and their gametes) to predominantly degraded (i.e., decomposing carcasses) within a single spawning season.

In addition to this well marked phenology, Pacific salmon systems have been a model system for decades, thus having relatively well-characterized molecular biology including transcriptional changes in steroidogenic pathways, gamete production, and reproductive behavior (Mommsen et al., 2004). Adult and juvenile fish exhibit distinct DNA methylation patterns associated with developmental maturation (Ruiz et al., 2025). Spawning aggregations concentrate specific pathogens including bacterial and fungal opportunists that proliferate on senescent tissue (Cipriano and Bullock, 2001; Van West, 2006). This biological knowledge enables hypothesis-driven environmental omics: we can predict when specific molecular signals should appear, test whether environmental samples capture expected temporal dynamics, and validate inferences against independent data sources like visual surveys and hatchery escapement records (Ip et al., 2025a).

Here we use shotgun Oxford Nanopore sequencing with adaptive sampling and modified basecalling to simultaneously recover environmental RNA, DNA methylation profiles, and pathogen signatures from water samples collected during Pacific salmon spawning runs. We evaluate how these molecular signals change over time, seeking to understand when they align, when they diverge, and what these reveal about the temporal boundaries of environmental inference. This raises two linked questions: what biological information can shotgun environmental sequencing recover beyond species presence, and when can each class of signal be trusted given that environmental molecules degrade at different rates? Specifically, we ask: (i) whether total environmental messenger RNA tracks spawning phenology; (ii) whether methylation-based demographic inferences are biologically meaningful in environmental samples; and (iii) whether eRNA, methylation, and pathogen signals each track spawning activity at timescales consistent with their environmental persistence.

Through systematic analysis of Coho and Chinook salmon spawning in Issaquah Creek, we reveal a short-lived temporal window, termed the’Freshness Gate’, where environmental nucleic acids most faithfully capture biological state as opposed to degraded signals (Figure 2). This finding establishes practical and theoretical boundaries for multiomic inference in natural ecosystems and provides a framework for designing environmental monitoring programs that account for molecular signal lifetimes, maximizing the biological information recoverable from each sample beyond simple species presence.

**Figure 2.**
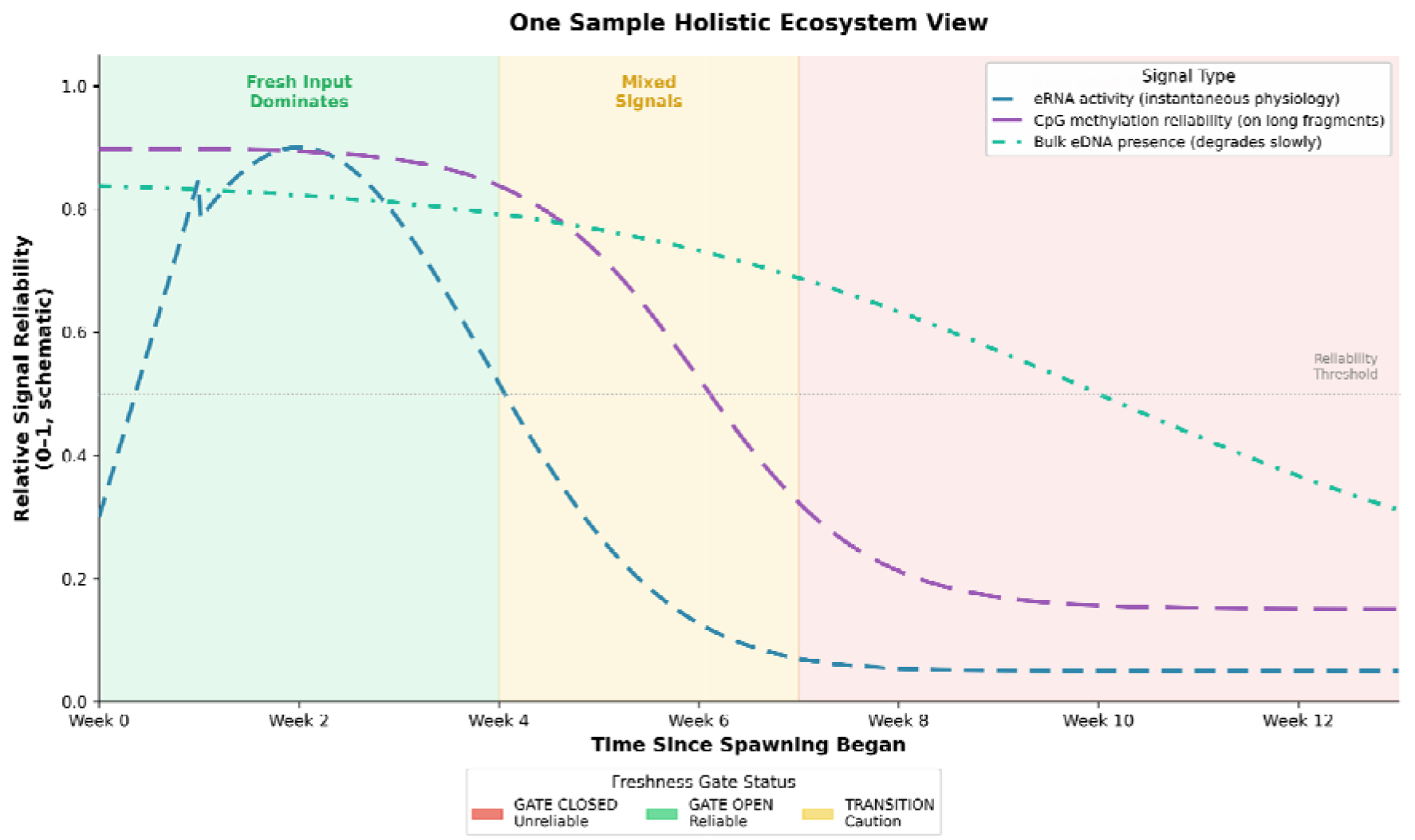
Conceptual model: Information reliability continuum across spawning season. Schematic (NOT empirical data) illustrating relative signal reliability from environmental nucleic acids over time. Three signal types show distinct temporal dynamics: environmental RNA activity (blue) provides short-lived physiological signals (hours-days); CpG methylation reliability (purple) yields demographic information only within a’Freshness Gate’ when fresh DNA dominates and long fragments are present; bulk eDNA presence (teal) persists longest, reflecting occurrence/abundance. Background shading denotes operational phases: green = GATE OPEN (reliable demographic inference, fresh DNA dominates); yellow = TRANSITION (mixed signals, caution required); red = GATE CLOSED (unreliable, degraded DNA dominates). Horizontal dashed line marks reliability threshold (0.5). Demographic inference requires both conditions: (i) fresh-input evidence (environmental RNA spawning spike or qPCR mtDNA elevation) and (ii) sufficient long-fragment fraction. Empirical validation of this framework is presented in Figures 3-4.

## 2. Methods

### 2.1 Study system and sampling design

We conducted weekly water sampling during the fall 2024 salmon spawning season at Issaquah Creek, a tributary of Lake Washington in western Washington State. Issaquah Creek supports natural runs of Coho (*Oncorhynchus kisutch* (Walbaum, 1792)) and Chinook (*O*. *tshawytscha* (Walbaum, 1792)) salmon with well-documented spawning phenology. The Issaquah Salmon Hatchery operates a fish ladder at which returning adults are counted, providing an independent record of run timing for both species. Our sampling location was positioned immediately downstream of the ladder gate, adjacent to natural aggregations in the creek channel (47.5295°N, 122.0391°W). Water samples were collected concurrently with those used for companion studies on airborne eDNA detection (Ip et al., 2025a) and air-water metabarcoding sampling (Ip et al., 2025c), with each 3 L collection split into three 1 L biological replicates allocated across studies; the present study used the third replicate for environmental omics analysis.

Sampling occurred weekly from September 18 through November 20, 2024 (designated Weeks 1 - 9), spanning the Chinook run, Coho arrival and peak spawning (late October through early November), and post-spawning carcass decomposition phases. At each sampling event, we collected three biological replicates of 1 liter surface water each, for a total of 3 liters per week. This design provided temporal coverage of the complete spawning season while maintaining consistent sampling effort and location to isolate temporal variation from spatial effects.

Hatchery escapement data documenting daily counts of Coho and Chinook salmon passing through the fish ladder were obtained from Washington Department of Fish and Wildlife (Supplementary Figure S8). These counts record fish entering the hatchery facility rather than fish spawning in the creek channel where sampling occurred, and were recorded at irregular intervals; they are therefore used as a qualitative record of run timing rather than as a quantitative abundance reference.

### 2.2 Sample preservation, nucleic acid extraction, reverse transcription

Water samples were filtered on-site using a Smith-Root eDNA Citizen Science Sampler equipped with 5.0 μm pore size mixed cellulose ester (MCE) filters. Each 1 liter replicate was filtered separately, and filters were immediately preserved in 1.5 mL of DNA/RNA Shield buffer (Zymo Research) using sterile disposable forceps. This preservation method stabilizes both DNA and RNA without requiring immediate freezing, enabling field processing at the creek site. Field-negative controls consisting of 1 liter Milli-Q water were processed using the same filtration system to assess contamination during sample collection and processing. All equipment was decontaminated between sampling events using 10% bleach solution followed by thorough rinsing with deionized water. Preserved filters were stored at-20°C and extracted as a single batch within one week of the final collection date, so early-season samples were stored for up to 9 weeks prior to extraction. Batch extraction removes processing date as a source of variation across the time series. Storage duration was inversely related to collection date, so any degradation introduced during storage would act most strongly on the earliest samples; because those samples show the freshest molecular signal, storage artifacts cannot account for the observed temporal pattern.

Field-negative controls were not sequenced in the present study due to barcode constraints on the nanopore multiplexing kit (SQK-NBD114-24). However, field blanks and PCR no-template controls collected concurrently with the same filtration equipment and processed through identical preservation workflows were sequenced and analyzed in two companion studies using the same biological replicates (Ip et al., 2025a; Ip et al., 2025c). In both studies, all negative controls showed no detectable salmon DNA by qPCR or metabarcoding, confirming negligible contamination during field collection.

We processed samples using two separate extraction workflows optimized for RNA versus DNA analyses. For environmental RNA samples, 300 μL of the buffer in which the filters were stored was thawed on ice and extracted using the Qiagen RNeasy Mini Kit following manufacturer protocols. RNA fractions were treated with DNase I (Qiagen) to remove residual DNA contamination, purified through RNeasy MinElute columns, and eluted in RNase-free water. RNA integrity and quantity were assessed using an Agilent 2100 Bioanalyzer. To enable robust detection and sequencing of low-abundance environmental transcripts, we performed Sequence-Independent Single-Primer Amplification (SISPA) adapted from Chrzastek et al. (2017). Briefly, 10 µL of total environmental RNA was reverse transcribed using random hexamer primers with SuperScript IV Reverse Transcriptase. First-strand cDNA was converted to double-stranded cDNA using Klenow fragment with Primer A (5’-GTTTCCCACTGGAGGATA-N9-3’), purified with AMPure XP beads, and amplified by PCR using Primer B (5’-GTTTCCCACTGGAGGATA-3’) with LongAmp Taq 2X Master Mix (30 cycles: 94°C/15s, 50°C/30s, 65°C/2min) to generate double-stranded cDNA. This universal amplification approach enriches all RNA templates without target-specific priming, effectively increasing RNA copy numbers across the entire transcriptome to facilitate downstream shotgun sequencing.

For DNA methylation samples, 300 μL of DNA/RNA Shield preservative was extracted directly using phenol-chloroform-isoamyl alcohol (PCI) following a modified protocol (Supplementary Material S1). Briefly, the preservative aliquot was digested with proteinase K (final concentration 1 mg/mL) at 56°C for 2 hours, extracted twice with phenol:chloroform:isoamyl alcohol (25:24:1) using phase-lock tubes, followed by chloroform:isoamyl alcohol (24:1) cleanup. DNA was precipitated with isopropanol at room temperature, pelleted by centrifugation at 13,300 × g for 30 minutes at 4°C, washed twice with ice-cold 70% ethanol, and resuspended in low-TE buffer at 37°C. This organic extraction method was selected to maximize recovery of high-molecular-weight genomic DNA with intact base modifications suitable for direct nanopore basecalling without bisulfite conversion. DNA was quantified using a Qubit fluorometer (Thermo Fisher) and quality-assessed by fragment length distribution on an Agilent 2100 Bioanalyzer.

### 2.3 qPCR quantification

Quantitative PCR assays provided an orthogonal measure of fresh biological input and enabled absolute quantification of salmon eDNA concentrations. We used published species-specific assays from Duda et al. (2021). For Coho salmon, we used the COCytb_980-1093 assay targeting mitochondrial cytochrome b, with primers COCytb_980-1093_F (CCTTGGTGGCGGATATACTTATCTTA) and COCytb_980-1093_R (GAACTAGGAAGATGGCGAAGTAGATC), amplifying a 114-bp fragment. For Chinook salmon, we used the CKCO3_464-534 assay targeting the mitochondrial cytochrome oxidase III / NADH dehydrogenase 3 (COIII/ND3) region, with primers ATTCCATGGCCTACACGTGA and TTGGTATTGGACCTGTCGCAGAAG, amplifying a 71-bp fragment; this assay is identical to that of Shelton et al. (2019) (Duda et al., 2021; Shelton et al., 2019). Both assays were run in SYBR Green format without the published TaqMan probes; specificity was therefore assessed by melt curve analysis. qPCR was initiated in Week 4 of the sampling series; no qPCR measurements are available for Weeks 1–3.

Reactions were performed using SYBR Select Master Mix (Fisher Scientific) on an Applied Biosystems QuantStudio 5 real-time PCR system with 384-well plates. Each 10 μL reaction contained 5 μL SYBR Select Master Mix, 0.4 μL forward primer (10 μM), 0.4 μL reverse primer (10 μM), 2.2 μL molecular-grade water, and 2 μL template DNA. Thermal cycling conditions were: 95°C for 10 min, followed by 40 cycles of 95°C for 15 sec and 60°C for 60 sec. Melt curve analysis confirmed amplification specificity, with positive standards showing a single assay-specific peak (79–81°C) and no-template controls showing no amplification.

Standard curves were constructed using tissue-extracted DNA from adult Coho and Chinook salmon quantified by Qubit fluorometry. Serial dilutions spanning 10 to 10¹ copies/μL were prepared and run in triplicate to quintuplicate depending on concentration. qPCR performance was characterized by amplification efficiency, detection probability across the standard curve range, and limit of detection with propagated uncertainty (Supplementary Figure S9). Environmental sample DNA concentrations were estimated using hierarchical Bayesian observation models that account for both technical qPCR variation and biological concentration uncertainty, following Guri et al. (2024) and Ip et al. (2025a). This probabilistic framework provides robust concentration estimates even near detection limits where binary presence/absence would be unreliable.

### 2.4 Oxford Nanopore library preparation and sequencing

Two separate sequencing libraries were prepared for environmental RNA (cDNA) and DNA methylation analyses using Oxford Nanopore Technologies (ONT) Native Barcoding Kit 24 V14 (SQK-NBD114-24). For eRNA libraries, 200 ng of SISPA-amplified cDNA per sample was used as input per manufacturer’s recommendations. For DNA methylation libraries, up to 200 ng of PCI-extracted genomic DNA per sample was used as input, with the full available yield loaded for low-concentration samples, with no amplification or size selection, preserving native modifications for direct basecalling. Each library was sequenced on two R10.4.1 flow cells, one with adaptive sampling enabled and one standard, for four runs total. Primary analyses used the adaptive sampling runs for both eRNA and methylation; standard runs provided non-enriched comparisons for adaptive sampling validation (Supplementary Figure S4).

Nanopore “adaptive sampling” was configured for both the native genomic DNA and eRNA/cDNA flow cells to enrich for salmonid sequences while retaining unbiased background reads. Reference panels included complete genome assemblies (*O*. *kisutch*: GCF_002021735.2, Okis_V2; *O*. *tshawytscha*: GCF_018296145.1, Otsh_v2.0) for the DNA sequencing and their corresponding RefSeq mRNA transcript sets obtained from NCBI for the eRNA/cDNA sequencing. “Adaptive sampling” means that during sequencing, reads mapping to these references were sequenced to completion while other reads were rejected early by the pores, increasing on-target recovery approximately 3-fold (Payne et al., 2021) without distorting the relative proportions of species assignments among salmonid reads (Supplementary Figure S4).

Libraries were loaded onto R10.4.1 flow cells and sequenced on a MinION MK1D for 40-42 hours per run. Post-sequencing basecalling was performed using Dorado v0.9.1 in Super-Accurate mode (model: dna_r10.4.1_e8.2_400bps_sup@v5.0.0) on a local workstation equipped with an NVIDIA RTX 4090 GPU. For DNA methylation libraries, each modification type was basecalled separately with the corresponding modification model: 5mCG_5hmCG and 5mC_5hmC, producing BAM files with per-read methylation tags (MM/ML fields). Barcode assignment was integrated into basecalling using the --kit-name SQK-NBD114-24 flag, and demultiplexing was performed with Dorado demux using --no-classify to split pre-assigned barcodes without reclassification. Sequencing runs yielded 11.7 Gb (eRNA/cDNA adaptive sampling), 12.9 Gb (eRNA/cDNA standard), 2.4 Gb (native DNA adaptive sampling), and 1.6 Gb (native DNA standard) of estimated bases.

### 2.5 Bioinformatic processing and quality control

Demultiplexed BAM files were aligned to reference genomes using Dorado aligner v0.9.1 with default parameters. Reads shorter than 200 bp or with mean quality scores below Q10 were discarded. Passed reads were classified using multiple approaches: (1) taxonomic assignment with Kraken2 (Wood et al., 2019) against the NCBI core_nt database (downloaded November 2024) for community composition and pathogen detection; (2) salmon-specific transcriptome mapping using Minimap2 (Li, 2018) for environmental RNA quantification; (3) whole-genome alignment and species confirmation using Minimap2; and (4) methylation calling from basecalling outputs for CpG profiling.

Read statistics including N50 length, quality scores, and yield were computed using Dorado basecalling summary outputs and custom R scripts (see Data Availability). Negative controls from companion studies using the same field collection pipeline showed no detectable salmon contamination (Ip et al., 2025a; Ip et al., 2025c), and independently prepared sequencing runs showed consistency in species detection and relative abundance patterns (Pearson correlation r > 0.85 across weeks).

### 2.6 Environmental RNA transcript detection and categorization

Transcript identification followed a reference-based mapping approach adapted for degraded environmental RNA. Reads were mapped to annotated transcriptomes for *Oncorhynchus kisutch* and *O*. *tshawytscha* using Minimap2 (-ax map-ont). Only reads mapping with MAPQ ≥ 20 and covering ≥ 80% of their length were retained. This stringent filtering reduces false-positive assignments arising from degraded fragments and from homologous genes shared between the two species.

SISPA involves non-specific whole-transcriptome amplification, so absolute transcript abundances are not preserved (Chrzastek et al., 2017; Djikeng et al., 2008). We therefore treat environmental RNA as semi-quantitative. Because all samples passed through an identical extraction, amplification, and sequencing workflow, we compare relative changes in category-level read proportions across time points rather than interpreting values as absolute expression levels. **A** gene was reported as detected when at least one read passed the mapping filters described above. Interpretation is accordingly anchored at the category level; individual genes recovered at fewer than approximately 50 reads across the season are treated as provisional detections rather than confident calls, and no conclusion in this study rests on such a gene alone.

Detected transcripts were assigned to five functional categories with established roles in salmon spawning physiology (Mommsen et al., 2004; Table 1): (1) Spawning (Reproductive), comprising extracellular matrix **component**s, sex determination genes, steroidogenic enzymes, and reproductive hormone receptors (Russell and Robker, 2007; Matson et al., 2011); (2) Death (Apoptosis/Senescence), comprising initiator caspases, caspase-associated regulators, and matrix metalloproteinases (Li et al., 1997; Salvesen and Dixit, 1997; Sternlicht and Werb, 2001); (3) Neuroendocrine (Spawning Behavior), comprising prolactin, oxytocin, and isotocin receptor transcripts (Hausmann et al., 1995); (4) Environmental Adaptation (Osmoregulation/Stress), comprising Na /K-ATPase subunits and interacting proteins (Blanco and Mercer, 1998), hypoxia response factors, and heat shock chaperones; and (5) Immune/Inflammatory, comprising anti-inflammatory cytokines and inflamma**tory** regulators.

**Table 1.**
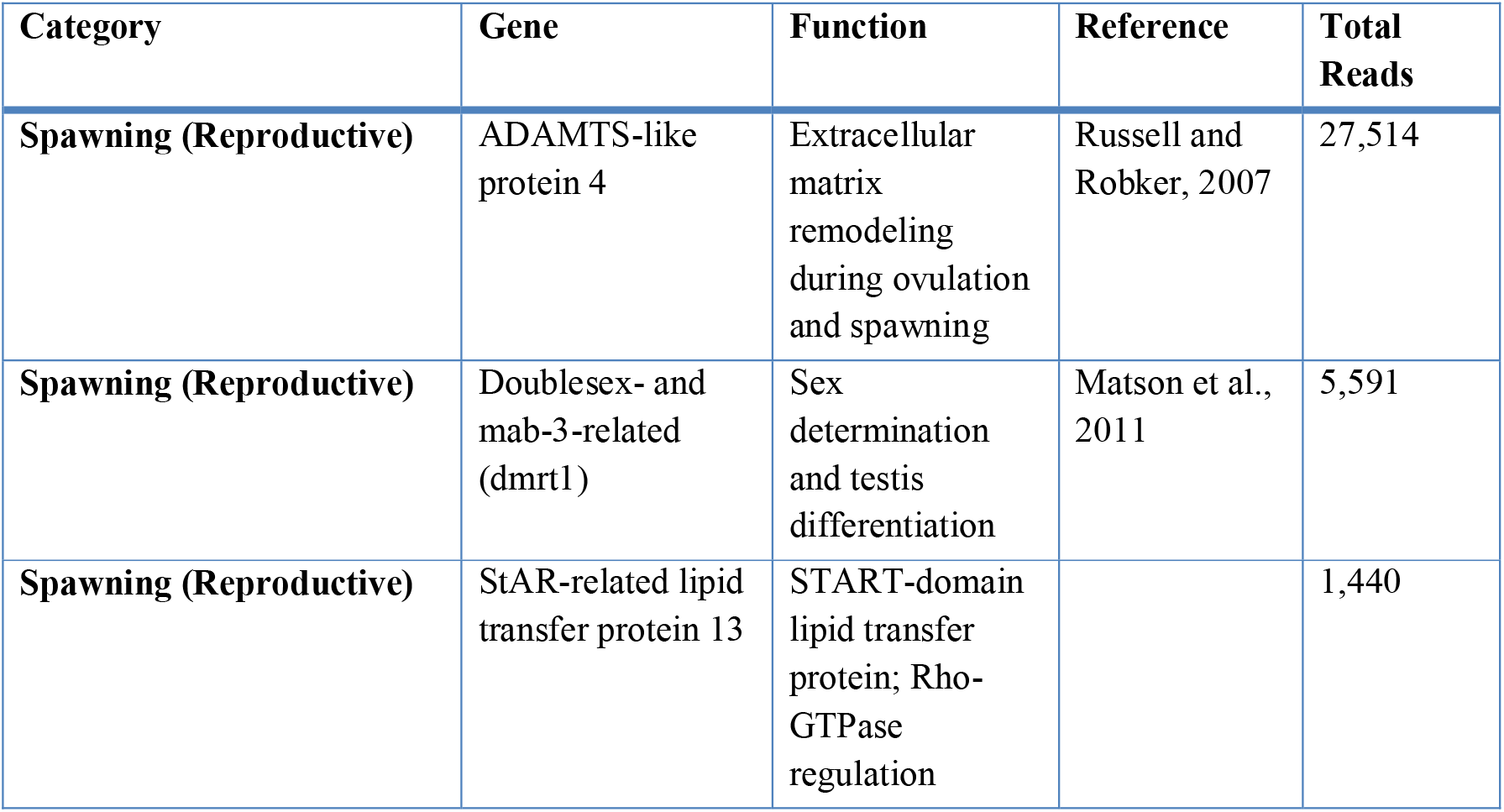
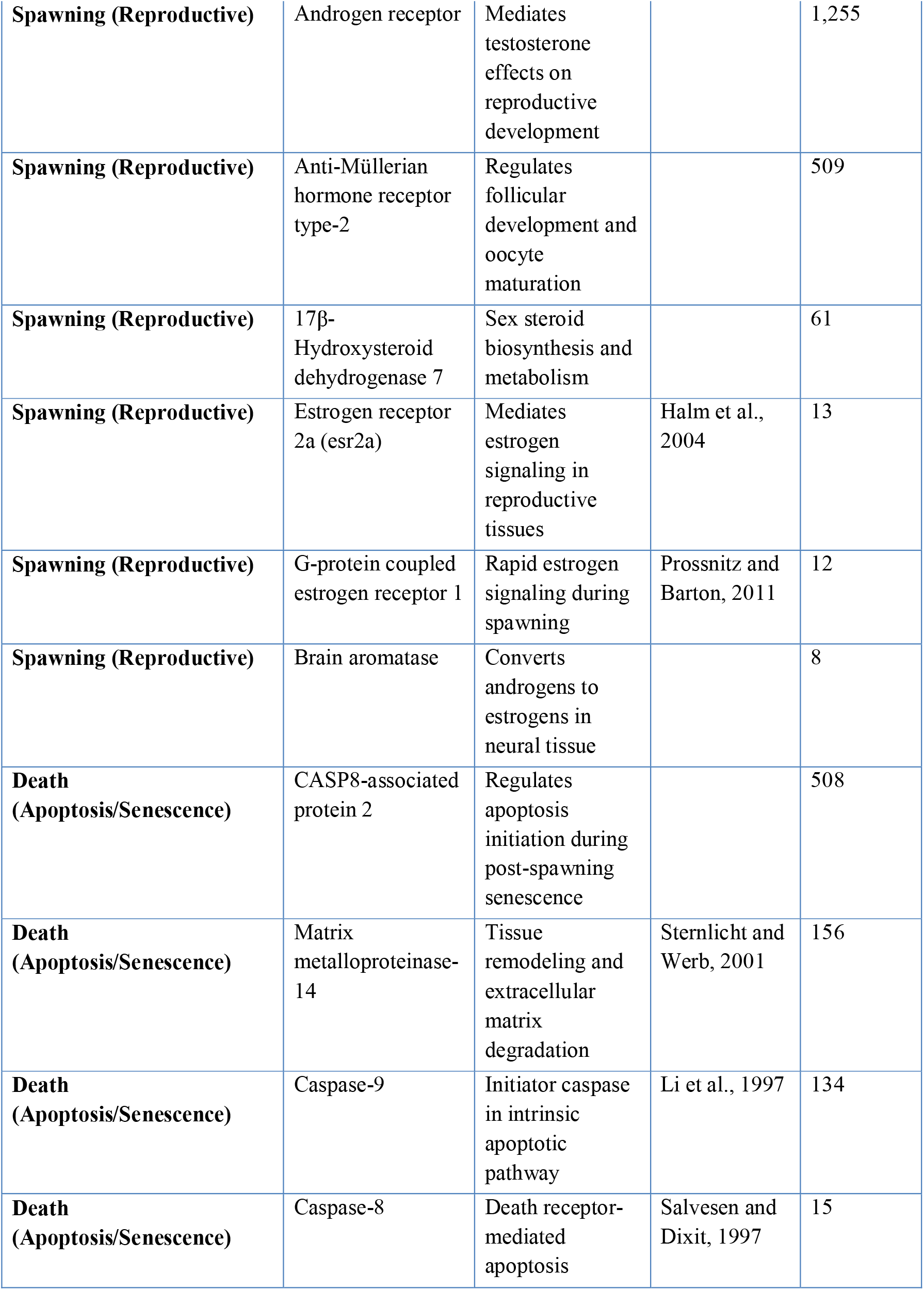
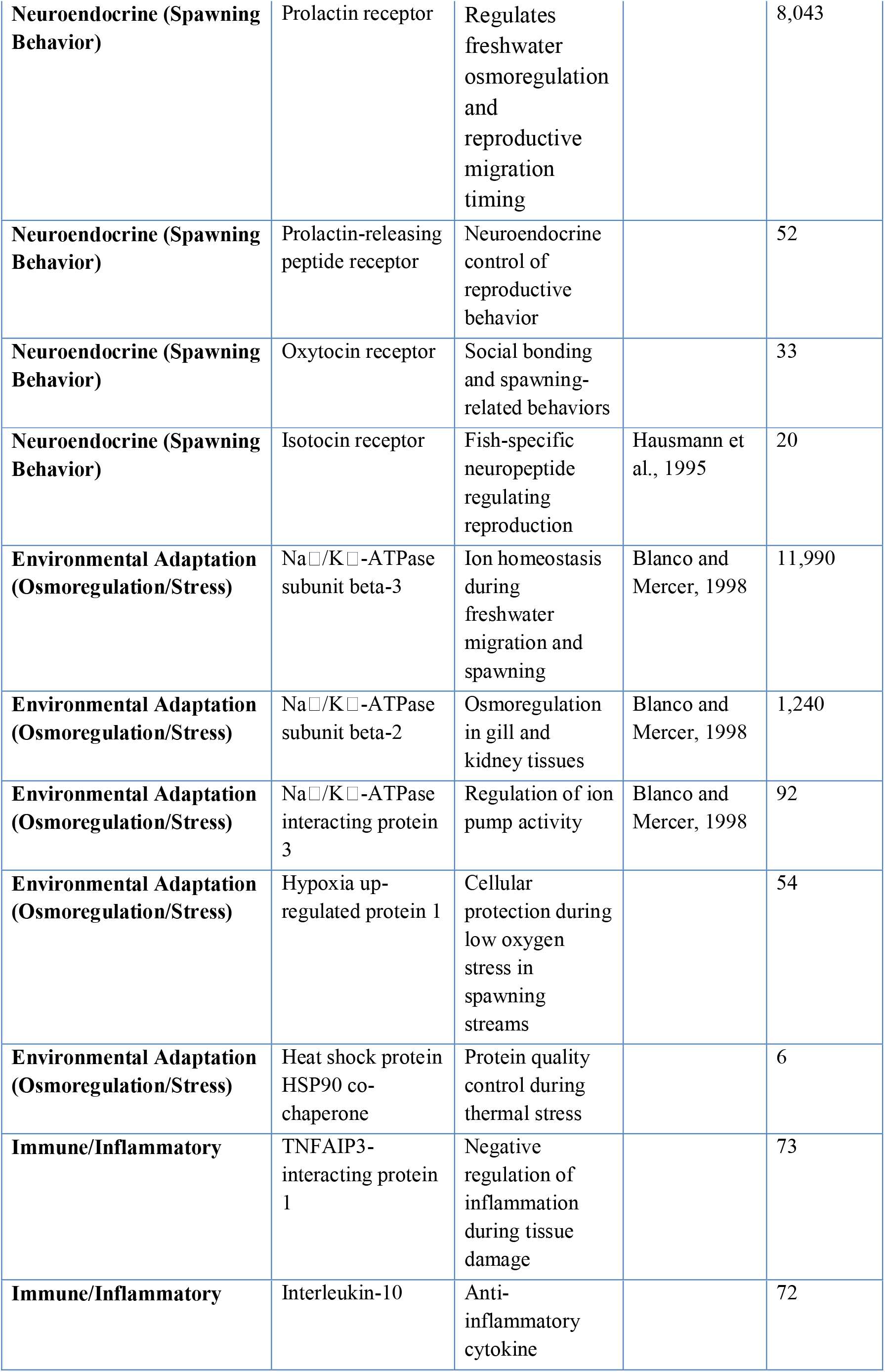
Representative transcripts detected from environmental RNA shotgun metatranscriptomics in Coho salmon. (*Oncorhynchus kisutch*). Genes shown represent the most abundant transcripts from each functional category across 9 weeks of environmental nucleic acid sampling (Weeks 1-9) using adaptive nanopore sequencing. Total reads represent cumulative detection across all field samples. Complete gene list provided in Supplementary Material S2.

Category assignment was performed by keyword matching against gene names in the reference annotation, followed by manual review of the highest-abundance genes in each category. Automated assignment of this kind is efficient at scale but can misclassify genes whose names share a domain family with functionally distinct members; category-level patterns are therefore more reliable than assignments for any individual gene.

Histone transcripts (H2A, H2B, H3, H4) were tracked separately from these five categories, as a proxy for the total quantity of salmon mRNA recovered in each sample rather than as a stable normalizing reference. Histone recovery varied substantially across the season, and that variation is treated as signal reflecting changing mRNA input rather than as technical noise to be normalized away (Figure 3B).

**Figure 3.**
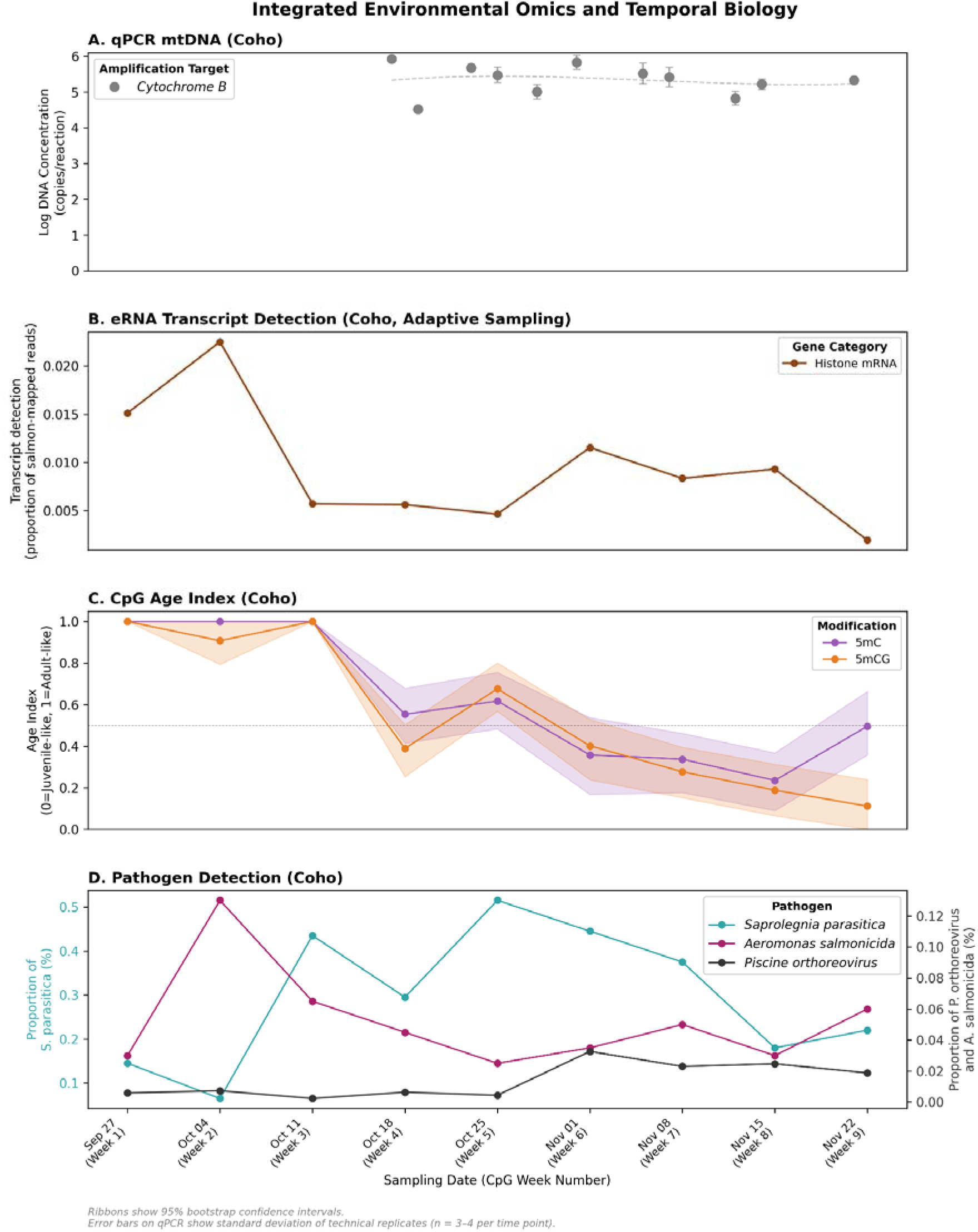
Integrated environmental omics reveal’Freshness Gate’ windows (Coho salmon). Multi-platform temporal signals from qPCR, environmental RNA, methylation, and pathogens define brief periods when environmental nucleic acids most accurately reflect living biological input. (A) qPCR mtDNA quantification (log10 DNA copies per reaction); error bars show standard deviation of technical replicates (n = 3-4 per time point). Dashed line in Panel A shows a trend fit for visual guidance only. qPCR sampling began in week 4, so no qPCR measurements are available within the earliest sampling weeks. (B) Histone mRNA recovery, expressed as the proportion of salmon-mapped reads, across nine weeks. Histone mRNA varied 11.5-fold across the season (CV = 66.7%), was independent of sequencing depth (r = 0.03, p = 0.94), and replicated across two independently prepared sequencing runs (r = 0.90, p = 0.001). The transcripts detected in each functional category are listed in Table 1. (C) Methylation (CpG) Age Index projected onto juvenile and adult references; ribbons show 95% bootstrap confidence intervals and the dashed line marks the 0.5 adult-like threshold. (D) Pathogen detection dynamics, as the percentage of total sequencing reads per library classified to each taxon by Kraken2. *Saprolegnia parasitica* (left y-axis) is plotted on a separate scale from *Aeromonas salmonicida* and *Piscine orthoreovirus* (right y-axis) due to the large difference in proportions. High *Saprolegnia* abundance (peak 51.5% at week 5) reflects oomycete colonization of decomposing salmon tissue during late-run phases.

Category-level signal was expressed as the proportion of total salmon-mapped reads assigned to each category, normalized by gene length to account for size bias (Wagner et al., 2012). Temporal dynamics were plotted on a log scale to resolve low-abundance categories alongside high-abundance ones. Gene identities, functional annotations, supporting references, and total read counts are provided in Table 1.

### 2.7 DNA methylation profiling and Age Index calculation

DNA methylation was profiled directly from native nanopore basecalling outputs without bisulfite conversion. Methylation calls were extracted from aligned BAM files using modkit v0.4.5 (https://github.com/nanoporetech/modkit) with the extract full command, which parses per-read modification tags (MM/ML fields) produced during Dorado’s modified basecalling. Per-site methylation fractions were computed as the ratio of methylated to total calls at each genomic position, and merged across samples into unified matrices—where each row represents a genomic CpG position and each column a weekly sample, with values representing the methylation fraction (methylated calls / total calls)—using custom Python scripts for matrix construction and filtering. Sites were retained through a two-stage filtering process: first, sites were required to be detected in both adult and juvenile calibration samples (regardless of methylation level) and in at least two of nine environmental eDNA samples, reducing the initial matrix from approximately 1.4 million to approximately 21,000 sites; second, for temporal analyses, sites were further required to be detected in at least five of nine weekly eDNA samples, yielding 1,555 sites for 5mC and 1,078 sites for 5mCG.

This progressive filtering ensures that retained sites have sufficient coverage for reliable methylation fraction estimation across the full time series; sites detected in fewer samples likely reflect stochastic sequencing depth variation rather than biological signal. While these age-informative loci represent a small fraction of the initial CpG sites, the approach parallels the selection of informative SNPs from genome-wide panels for population assignment: a small number of loci with high discriminatory power is sufficient for reliable classification. Sensitivity analyses using relaxed filtering thresholds (2-of-9 eDNA samples) retained more sites but yielded consistent Age Index trajectories, confirming that the stringent filter improves precision without altering biological conclusions. Tissue samples for adult calibration consisted of muscle tissue from wild-caught adult Coho and Chinook salmon (age 3+ years) purchased from Pike Place Market (Seattle, WA). Juvenile calibration samples consisted of tank water collected from juvenile Coho and Chinook rearing tanks (<1 year old) at the Seattle Aquarium in December 2024. Species identity for all calibration samples was confirmed by qPCR using the primers described in Section 2.3.

Adult tissue DNA was extracted using PCI (Section 2.2) to preserve methylation; juvenile tank water was filtered and preserved identically to environmental samples before PCI extraction. We note that adult calibration represents tissue-derived DNA while juvenile calibration represents waterborne eDNA, paralleling the environmental samples more closely for the juvenile calibration endpoint.

Age-informative CpG sites were identified by comparing methylation fractions between adult and juvenile salmon tissue calibration samples. Specifically, we selected CpG sites showing |adult - juvenile methylation| > 1% (i.e., absolute difference ≥ 0.01), which represent loci undergoing developmental methylation or demethylation during maturation. This threshold yielded 455 age-informative sites for 5mC and 364 sites for 5mCG in Coho, and 1,100 sites for 5mC and 714 sites for 5mCG in Chinook (Supplementary Figure S3).

The Methylation Age Index was computed as an effect-size-weighted projection onto the adult–juvenile axis defined by tissue calibration samples. Each age-informative CpG site was weighted by its absolute effect size (|adult methylation − juvenile methylation|), so sites with larger developmental differences contribute proportionally more to the index. Formally, for each week t and modification type m:

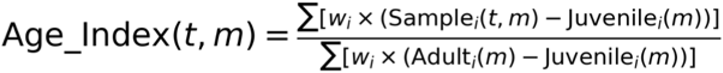

where w_i = |Adult_i(m) − Juvenile_i(m)| and the summation runs over all age-informative CpG sites. This transformation scales methylation fractions to a 0-1 index where 0 indicates juvenile-like methylation patterns, 1 indicates adult-like patterns, and intermediate values reflect mixtures or biological variation. An Age Index of 1.0 means the sample’s methylation pattern matches the adult tissue reference, not that all CpG sites are 100% methylated; for example, adult Coho tissue shows raw methylation fractions of approximately 56.6% for 5mC and 24.7% for 5mCG (Supplementary Figure S3). An R implementation of the Age Index calculation is provided in the accompanying repository to facilitate application to other systems.

Uncertainty in the Age Index was estimated using bootstrap resampling (1000 iterations) across CpG sites, generating 95% confidence intervals for each week. Age Index values above 0.5 were operationally defined as adult-like, consistent with the expectation that spawning-run samples should be dominated by adult-origin DNA when fresh input is high. Fragment-length sensitivity analyses (Supplementary Figure S2) indicate that Age Index patterns are consistent across the 1-2 kb and 2-5 kb size fractions in Coho, while the 0-500 bp fraction and the Chinook samples show greater variability.

Three assumptions underlie this Age Index formulation. First, the adult-juvenile methylation axis is defined by a single calibration sample per life stage (adult tissue from one wild-caught individual per species; juvenile eDNA from one rearing-tank collection), so the index captures a relative ordering along that axis rather than an absolute demographic assignment; biological replication across individuals and populations would strengthen the calibration (see Limitations). Second, the weighted projection assumes a linear relationship between methylation fraction and demographic composition at each CpG site, without modeling intermediate life stages (smolts, sub-adults) that may show non-linear methylation trajectories. Third, weighting by |delta| gives greater influence to sites with larger adult-juvenile differences, analogous to FST-weighted panels in population genetics or loading-weighted projections in PCA; this is appropriate when the goal is maximizing discrimination between endpoints but could overweight sites whose large differences reflect tissue-specific rather than age-specific biology, given the asymmetry between adult tissue-derived and juvenile waterborne calibration DNA.

### 2.8 Pathogen detection and community profiling

Taxonomic classification was performed on simplex-basecalled (unmodified) reads from both eRNA/cDNA sequencing runs (ONT006, standard sequencing; ONT007, adaptive sampling) using Kraken2 v2.1.2 (Wood et al., 2019). Reads were classified in two independent passes: first against the MitoFish mitochondrial reference database for vertebrate species assignment, and second against the NCBI core_nt database (downloaded November 2024) for community composition and pathogen detection. Classification used Kraken2’s default confidence threshold (0.0), a permissive setting that maximizes sensitivity for low-abundance taxa but increases spurious assignment, which motivated the alignment-based validation described below. The core_nt database includes bacterial, fungal, viral, and eukaryotic genomes enabling broad-spectrum pathogen and community detection. Reads assigned to known salmon pathogens were extracted for temporal analysis. We focused on three pathogen classes representing distinct transmission ecologies and identified as management priorities through consultation with Issaquah Salmon Hatchery staff (Krkosek et al., 2024; Bass et al., 2023): *Aeromonas salmonicida* (bacterial furunculosis; 10,625 total reads), *Saprolegnia parasitica* (oomycete saprolegniasis; 2,187 reads), and *Piscine orthoreovirus* (PRV, viral disease common in Pacific salmon; 2,313 reads). Additional known salmon pathogens were also detected, including *Flavobacterium psychrophilum* (bacterial cold water disease; 20,140 reads, the most abundant pathogen detected in the dataset), *Ceratonova shast*a (myxozoan ceratomyxosis; 9,963 reads), and *Tetracapsuloides bryosalmonae* (proliferative kidney disease; 130 reads), though detailed temporal analysis of these taxa was beyond the scope of this study. We note that pathogen classification via Kraken2 k-mer matching provides rapid taxonomic assignment but does not confirm viability or infection status; these detections indicate the presence of pathogen-derived nucleic acids in the environmental water samples.

Kraken2 pathogen assignments were independently validated by minimap2 (Li, 2018) alignment of demultiplexed reads to reference genomes for the two focal bacterial and oomycete taxa (*A. salmonicida* NC_009348.1; *S. parasitica* ITS/actin markers), for PRV-1 genome segments, and for the most abundant detected pathogen (*F. psychrophilum* NC_009613.3). Genome-wide read coverage consistent with true pathogen presence was confirmed for *F. psychrophilum*, *A. salmonicida*, and *S. parasitica* (Supplementary Figure S12). PRV-1 showed no aligned reads despite Kraken2 classification, suggesting that PRV k-mer matches may reflect shared sequence features with related viral taxa, consistent with the permissive confidence setting, rather than true PRV presence; PRV detections should therefore be interpreted cautiously. Validation scripts and coverage data are available in the accompanying repository (https://github.com/piedna/Wild-Omics).

Pathogen abundance was expressed as the percentage of total library reads classified to each taxon. Because adaptive sampling rejects reads not matching the salmonid reference panel, non-target proportions differ between the two runs; pathogen abundances are therefore reported as relative enrichment within each library rather than as absolute community composition. Background community composition (bacteria, fungi, eukaryotes) was retained for context but was not the primary focus of pathogen analysis.

Cross-correlation analysis between pathogen abundances and salmon mRNA signals quantified temporal offsets (see Section 2.9 for statistical details).

### 2.9 Statistical analyses and variance partitioning

Cross-correlation analyses quantified temporal relationships between environmental mRNA signals (transcript detection as a proxy for total salmon mRNA input) and methylation Age Index. We compared the weekly mRNA signal against the Age Index at two offsets: the same sampling week, and with the mRNA signal shifted one week earlier, corresponding to mRNA leading methylation. Associations at each offset were tested using Pearson’s product-moment correlation. This analysis tests whether mRNA detection, which marks immediate biological input, temporally leads or lags the methylation Age Index, which integrates fresh and degraded DNA sources over longer timescales. Pathogen abundances were compared against the mRNA signal across offsets of zero to three weeks using the same approach, with the reported lag corresponding to the offset of maximum correlation.

PERMANOVA (Permutational Multivariate Analysis of Variance) tested the relative importance of biological versus technical factors (fragment length and read depth) in structuring methylation patterns. The response variable was a Bray-Curtis dissimilarity matrix computed from site-level methylation fractions across all age-informative CpG sites. Predictor variables included: (1) Age Index as a biological freshness gradient; (2) Fragment Length (median per week) representing physical degradation; and (3) Read Depth (total coverage) reflecting sequencing effort and DNA concentration. We note that Read Depth correlates with biological DNA abundance rather than pure technical variation, as high-abundance weeks (fish present, active spawning) naturally yield more DNA and thus more reads. Partial R² values quantified unique variance explained by each predictor after accounting for covariation. Because the Age Index is derived from the same methylation fractions used to construct the dissimilarity matrix, its partial R² is not independent of the response; this comparison is therefore interpreted as a test of whether technical covariates rival biological structure, not as an unbiased variance decomposition.

Principal Coordinates Analysis (PCoA) provided complementary multivariate assessment of Age Index trajectories. We computed Bray-Curtis dissimilarities among weeks based on methylation profiles, performed PCoA ordination, and plotted samples in the first two principal coordinate axes alongside adult and juvenile calibration samples. Ordinations were performed separately for 5mC and 5mCG modification types to test consistency across modification types (Figure 4, Supplementary Figure S7). To confirm that ordination results were not sensitive to the choice of dissimilarity metric, we also computed Aitchison (compositional) distances from centered log-ratio transformed methylation fractions and assessed concordance between Bray-Curtis and Aitchison distance matrices using Mantel tests (Pearson correlation, 999 permutations).

**Figure 4.**
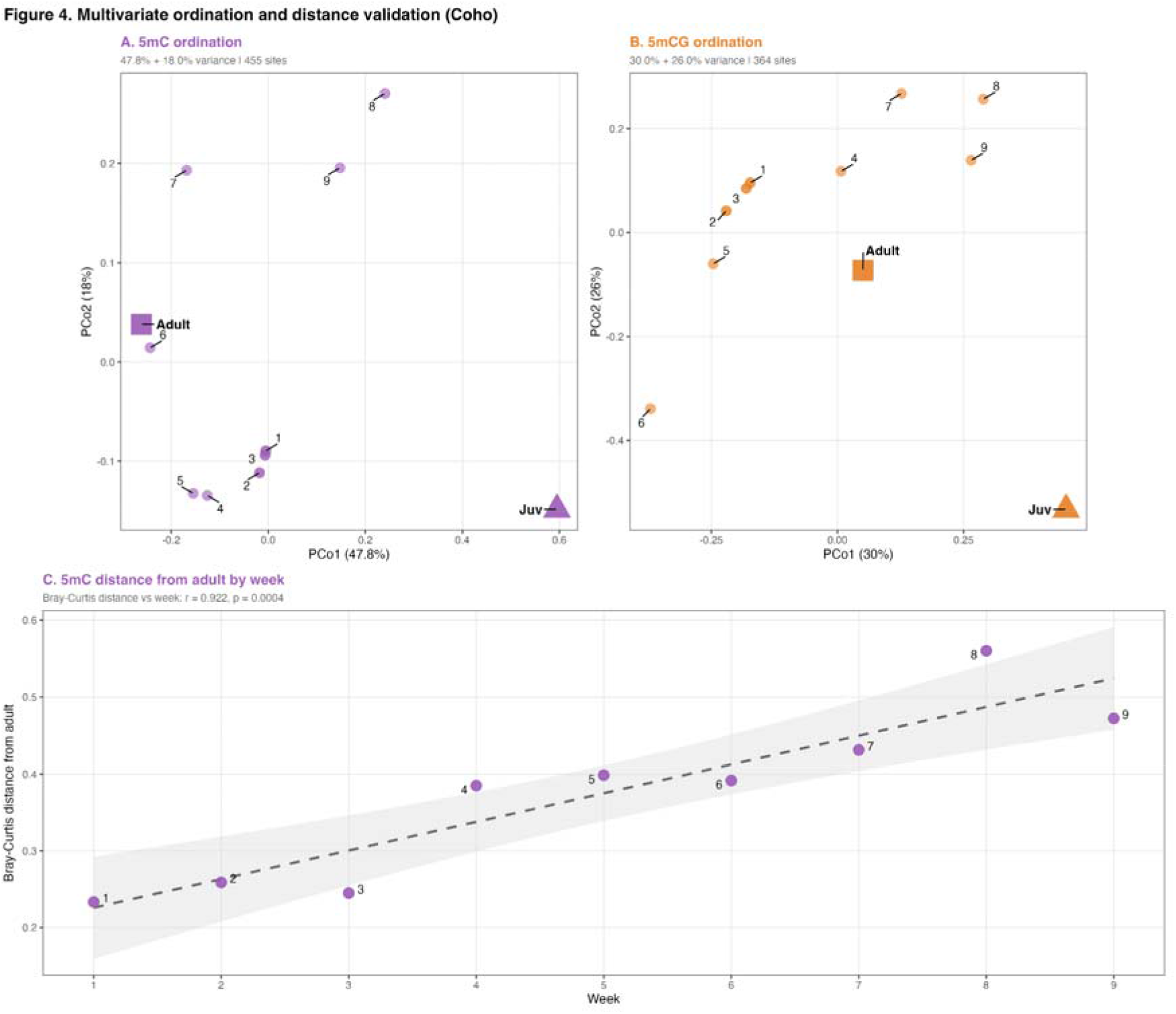
Multivariate ordination of age-informative CpG methylation patterns in Coho salmon. (A) 5mC ordination (Principal Coordinates Analysis) of age-informative CpG sites using Bray-Curtis distance: fresh-input weeks (1-3) cluster near the adult calibration reference (square), while degraded weeks (4-9) move progressively away from the adult reference, not specifically toward the juvenile reference, consistent with progressive loss of adult methylation. (B) 5mCG ordination shows the same departure with greater dispersion. (C) Bray-Curtis distance of each weekly sample from the adult calibration sample for 5mC increases progressively across the run (r = 0.922, p = 0.0004), a direct view of the same demethylation trajectory. Dashed line shows linear regression; grey shading indicates the 95% confidence interval. Because panels A and B are a two-dimensional projection of roughly two-thirds of the total variance while panel C uses the full distance matrix, proximity in the projection and the distances in C need not agree. Calibration samples: adult tissue, squares; juvenile eDNA, triangles. Selected week numbers (1, 3, 5, 9) label temporal progression.

All statistical analyses were conducted in R v4.3.2 (R Core Team, 2023) using packages vegan v2.6-4 (Oksanen et al., 2022) for community ecology and multivariate statistics, ggplot2 v3.4.0 for visualization, and custom scripts for methylation processing and pathogen validation in Python v3.8. Sequence alignment used minimap2 v2.26 (Li, 2018).

Significance threshold was set at α = 0.05 for all tests, though we emphasize effect sizes and confidence intervals over binary significance given the limited temporal resolution (n = 9 weeks) of our sampling design.

## 3. Results

### 3.1 Single-sample multi-omic readout of ecosystem state

Using shotgun Oxford Nanopore sequencing with adaptive sampling and modified basecalling, we simultaneously recovered environmental RNA, DNA methylation profiles, and pathogen signatures from individual water samples (Figure 1). Adaptive sampling increased on-target salmonid read recovery approximately 3.2-fold compared to non-enriched libraries (Supplementary Figure S4) while retaining background community reads for ecological context, enabling species-specific inference from the same library. This non-invasive workflow yields three concurrent biological readouts—mRNA detection, demographic inference, and pathogen signals—all from the same water sample.

### 3.2 Environmental mRNA detection and temporal dynamics

Across nine sampling weeks, Coho salmon environmental mRNA showed concordant temporal dynamics across all functional gene categories (Figure 3B, Table 1). All five functional categories, together with histone transcripts, peaked during the earliest sampling weeks (Weeks 1-2), declined through Weeks 3-5, and showed a secondary elevation during Coho arrival (Weeks 6-8) before falling to their lowest levels at Week 9. Read counts across detected transcripts spanned nearly four orders of magnitude (Table 1), reflecting stochastic variation inherent to shotgun recovery from complex environmental samples where detection probability depends on total salmon mRNA persistence, the proportion of salmon-derived reads relative to microbial background, and sampling depth.

All gene categories, together with histone transcripts, rose and fell in concert across the sampling period, indicating that these patterns reflect changes in total salmon mRNA abundance in the water column rather than differential gene expression (Figure 3B).

Environmental water integrates mRNA shed from multiple individuals, tissues, and degradation states; detecting a transcript in this medium reports its presence as biological input, not its regulation within any organism. Non-specific SISPA amplification further precludes quantitative comparison across transcript categories. Rather than resolving which genes are upregulated or downregulated, environmental mRNA detection reflects whether salmon-derived transcripts of any category are present in the water at detectable levels.

Temporal patterns in mRNA detection corresponded with known spawning activity. Transcript detection peaked during weeks of active salmon returns, with secondary elevations during later weeks (Weeks 6-8) coinciding with the arrival of late-season spawners (Supplementary Figure S8). The transcript signal declined during post-spawning weeks when carcass decomposition dominated and live fish were no longer contributing fresh mRNA to the water column.

Functional transcripts spanning all five categories were detected (Table 1), with their concordant temporal dynamics supporting the interpretation that detection reflects total mRNA input rather than category-specific biology. This temporal correspondence between mRNA presence and spawning activity establishes environmental RNA as a marker of fresh biological input, a property we formalize below as the Freshness Gate.

### 3.3 Convergent molecular signals define operational’Gate Open’ windows

Salmon environmental mRNA detection and qPCR mtDNA quantification tracked together over the weeks where both were measured, with Coho salmon mtDNA concentrations varying by approximately 1.4 log units across weeks 4–9 and peaking concurrently with mRNA detection (Figure 3A). qPCR sampling commenced in week 4, so mtDNA measurements are not available for weeks 1–3. However, these two markers report fundamentally different information: environmental RNA, with its hours-to-days persistence, indicates active fresh biological input, while qPCR DNA abundance reflects cumulative nucleic acid load including both fresh and degraded molecules. Their convergence identifies periods when fresh input dominates the environmental DNA pool, establishing operational’Gate Open’ windows for downstream methylation and demographic inference.

Specifically, weeks showing both elevated salmon mRNA (spawning-category reads > 0.1% of salmon-mapped reads) and qPCR mtDNA (> 10 copies/reaction; operational thresholds defined from natural breaks in the observed data distribution rather than independent calibration) were classified as Gate Open. For Coho salmon, Weeks 5 and 8 met this two-signal criterion. Weeks 1-3 were classified as Gate Open on elevated mRNA alone, since qPCR was not conducted in those weeks; this assignment is independently supported by the spawning pulses recorded in hatchery escapement data (Supplementary Figure S8).

Weeks with elevated qPCR but declining mRNA (e.g., Week 4) indicate persistent DNA without fresh input, which are precisely the conditions where methylation inference becomes unreliable.

### 3.4 Methylation demographics track fresh input within a reliability continuum

DNA methylation profiles projected onto adult-juvenile calibration references revealed that Age Index reliability exists on a continuum determined by the balance between fresh biological input and accumulated degraded DNA (Figure 3C). During weeks of high fresh input (Weeks 1-3), both 5mC and 5mCG Age Indices remained adult-like (median 5mC = 0.89, 5mCG = 0.91) with narrow bootstrap confidence intervals, indicating stable demographic inference (Supplementary Figure S1). However, during later Gate Open weeks when fish abundance had declined and environmental RNA indicated continued but lower spawning activity (Week 5), Age Indices remained adult-like but shifted toward intermediate values (5mC = 0.62, 5mCG = 0.68) with wider confidence intervals. This pattern is consistent with progressive dilution of fresh adult DNA by accumulated degraded material that has likely lost methylation marks.

This pattern reveals that methylation-based demographic inference requires sufficient concentration of fresh DNA to dominate degraded background signals, consistent with the Freshness Gate described above. The Gate is not a binary on/off switch but rather a reliability continuum where inference quality degrades progressively as the fresh:degraded ratio declines. Tissue calibration samples (Supplementary Figure S3) confirmed clear adult-juvenile separation, with adults more heavily methylated than juveniles (5mC: 0.566 vs. 0.260; 5mCG: 0.247 vs. 0.056). Because methylation increases with maturation, loss of methylation marks during environmental degradation moves adult-derived DNA toward juvenile-like values on this axis, the direction of drift observed in later weeks. By Week 8, despite meeting the Gate Open criterion, Age Indices had fallen below the adult-like threshold (Figure 3C), indicating that convergent mRNA and qPCR elevation is necessary but not sufficient for reliable demographic inference once accumulated degraded DNA dominates the pool.

### 3.5 Pathogens couple to host biology with ecological offsets

Shotgun metagenomic reads mapping to known salmon pathogens showed temporal dynamics coupled to host spawning biology with ecologically interpretable lags (Figure 3D). *Aeromonas salmonicida* (bacterial pathogen) tracked spawning activity with a cross-correlation peak at lag = +1 week (r = 0.68), though the peaks appear near-synchronous during early spawning weeks, consistent with rapid opportunistic bacterial proliferation following tissue damage and immune suppression during spawning stress. *Saprolegnia parasitica* (oomycete pathogen) exhibited more delayed peaks (+2 to +3 weeks relative to spawning maxima), reflecting saprotroph colonization of decomposing carcasses rather than infection of live fish. By mid-season (Week 5), *Saprolegnia* reads comprised 51.5% of total library content (including non-salmon reads), reflecting massive oomycete biomass during carcass decomposition.

Kraken2 classified reads as *Piscine orthoreovirus* (PRV) transiently in mid-season samples (Weeks 4-6); however, minimap2 alignment to PRV-1 genome segments produced no confirmed read coverage (Supplementary Figure S12), suggesting these classifications may reflect shared k-mer sequences rather than true PRV presence.

Together, the two confirmed pathogen signals, a bacterial opportunist and an oomycete saprotroph, display distinct temporal offsets relative to host spawning activity, each consistent with their known transmission ecology. Additional salmon pathogens detected but not selected as management-priority focal taxa, including *Flavobacterium psychrophilum* (bacterial cold water disease), *Ceratonova shasta* (ceratomyxosis), and *Tetracapsuloides bryosalmonae* (proliferative kidney disease), are consistent with the endemic pathogen community of Pacific Northwest freshwater systems, though their temporal dynamics were not analyzed in detail here.

### 3.6 Environmental mRNA leads methylation

Cross-correlation between the weekly histone mRNA signal (transcript detection serving as a proxy for total salmon mRNA input; Figure 3B) and the CpG methylation Age Index showed the following pattern (same-week correlations are positive but weak, 5mC r = 0.44, p = 0.241 and 5mCG r = 0.50, p = 0.169, n = 9; at a one-week mRNA lead the correlations strengthen, reaching significance for 5mC, r = 0.72, p = 0.043, and approaching significance for 5mCG, r = 0.65, p = 0.082, n = 8; Supplementary Figure S10). The 5mCG correlation, while not reaching the 0.05 threshold, is directionally consistent and of moderate effect size; given the limited sample size (n = 8 lagged pairs from 9 weekly samples), we emphasize effect sizes over binary significance thresholds. Because sampling occurred at weekly resolution, this offset represents a minimum detectable lag; the true biological coupling may occur over shorter timescales (days) that our sampling design cannot resolve. Nevertheless, the consistent one-week lead of mRNA over methylation across both modification types matches with transcript input marking immediate biological input, while methylation state reflects cumulative DNA composition that integrates fresh and degraded sources over longer timescales.

### 3.7 Biological freshness outweighs technical factors in methylation structure

PERMANOVA variance partitioning tested whether biological state (Age Index) or technical factors (Fragment Length, Read Depth) primarily organize environmental methylation patterns (Supplementary Figure S10E). Using Bray-Curtis dissimilarities of genome-wide CpG methylation as the response matrix, we found that Age Index explained the most unique variance for both modification types (partial R²: 5mC 0.131, 5mCG 0.251), above Read Depth (5mC 0.114, 5mCG 0.153) and Fragment Length (5mC 0.066, 5mCG 0.058). Although tests did not reach statistical significance (all p ≥ 0.05; n = 9 weeks, residual df = 5), effect magnitudes and directions are consistent with biological freshness gradients, rather than technical degradation metrics, driving methylation structure. These results are robust to the choice of dissimilarity metric, where Aitchison (compositional) and Bray-Curtis distance matrices correlate at r = 0.917 for 5mC and r = 0.859 for 5mCG (Mantel test, p = 0.001 for both, 999 permutations), and the PERMANOVA results are unchanged.

Read Depth correlates with biological DNA concentration rather than representing pure technical variation: high-abundance weeks (1-3, when fish are present) yield more DNA and deeper coverage, while low-abundance weeks (4+) yield less. Thus, Read Depth tracks the same fresh-input gradient as Age Index. Fragment Length, entered as the weekly median, explains the least variance, indicating that week-to-week differences in overall fragmentation do not drive methylation structure. Within individual samples, however, the age signal is most stable in fragments above 1 kb (Supplementary Figure S2), so demographic inference is best restricted to longer fragments even though sample-level fragmentation is not itself a major structuring factor.

### 3.8 Multivariate ordination corroborates Age Index trajectories

Principal Coordinates Analysis (PCoA) of age-informative CpG sites provided complementary support for Age Index patterns (Figure 4). In both 5mC and 5mCG datasets, fresh-input weeks (1-3) clustered tightly near adult tissue calibration samples in ordination space. In 5mC, later weeks (4-9) drifted progressively away from adult calibration samples; in 5mCG, later weeks showed greater dispersion without a clear directional trend, consistent with noisier modification calling at CpG-context sites. This ordination-based trajectory mirrors the Age Index time series (Figure 3C) but does not rely on the projection calculation, suggesting that temporal demethylation patterns are robust to methodological choices. The consistent fresh-phase clustering across both modification types corroborates the Age Index pattern independently of the projection calculation, though the stronger directional drift in 5mC suggests this modification may be more sensitive to progressive environmental degradation of methylation patterns. The same trend appears directly as Bray-Curtis distance from the adult calibration sample, which increases progressively across weeks for 5mC (Figure 4C; r = 0.922, p = 0.0004), whereas the 5mCG distance shows dispersion rather than directional drift (Supplementary Figure S11; r = 0.438, p = 0.238). Because the ordination in Figure 4A and B is a two-dimensional projection of roughly two-thirds of the variance whereas panel C uses the full distance matrix, proximity in the projection and distance in panel C need not agree.

### 3.9 Cross-species reproducibility with phenological offsets

Applying the multi-omic framework to Chinook salmon (*O*. *tshawytscha*) reproduced the Freshness Gate pattern with species-specific phenological timing (Supplementary Figures S5-S7). Cross-correlation analysis supports the mRNA-leads-methylation pattern in Chinook as well (5mC: r = 0.68 at lag =-1 week; 5mCG: r = 0.88 at lag =-1 week; Supplementary Figure S6), with Chinook showing stronger lag-1 correlations than Coho for both modification types, supporting cross-species robustness of the temporal coupling between transcript input and methylation dynamics. However, Chinook methylation showed greater heterogeneity and broader ordination dispersion compared to Coho. This reflects a data coverage constraint: Chinook environmental RNA was unavailable across a month-long window early in the series (Supplementary Figure S6), leaving fewer paired timepoints spanning the fresh-input phase for this species. In contrast, Coho sampling coincided with their run onset, enabling observation of the full temporal arc from fresh (Weeks 1-3) to degraded (Weeks 4+) states. Because histone genes are highly conserved across *Oncorhynchus* species, a fraction of reads assigned to one species’ histone reference may originate from the other; this potential cross-mapping does not affect temporal dynamics within a species but should be considered when comparing absolute histone mRNA proportions between Coho and Chinook.

This demonstrates that Freshness Gate inference depends on having sampling coverage across the fresh-input phase of a run. The apparent’failure’ of Chinook methylation inference actually serves as a negative control, confirming that degraded-phase sampling (analogous to Coho Weeks 6, 7, 9) yields unreliable demographics regardless of species.

Histone transcripts showed comparable temporal variation in Chinook (CV = 77.3%) and Coho (CV = 66.7%). Adaptive sampling improved species assignment purity comparably across both salmon species (Supplementary Figure S4), validating technical reproducibility of the environmental omics approach.

## 4. Discussion

### 4.1 From presence to process: Environmental omics as integrative ecosystem sensors

Environmental molecular monitoring has been dominated by amplification-based eDNA approaches optimized for answering a single question: which species are present (Thomsen & Willerslev, 2015; Ruppert et al., 2019)? While taxonomic inventories have transformed biodiversity assessment (Taberlet et al., 2018), they represent only a fraction of the biological information encoded in environmental nucleic acids (Cristescu & Hebert, 2018). The fundamental limitation has been both technological and conceptual. Long-read sequencing with native base modification detection, non-specific whole-transcriptome amplification, and adaptive sampling now provide the necessary tools. Yet, most environmental DNA studies continue to treat molecules as binary presence signals rather than information-rich biological records.

Our results demonstrate that parallel sequencing of environmental nucleic acids on a single platform converts environmental samples into multi-dimensional biosensors. Non-specific whole-transcriptome amplification (SISPA) of the RNA fraction captures biological input from environmental mRNA (Cristescu, 2019; Yates et al., 2021), while native DNA sequencing with base modification detection recovers age demographics from CpG methylation patterns (Balard et al., 2024; Horvath, 2013) and health from pathogen signatures (Bass et al., 2015) simultaneously. This integrated view, which we obtained non-invasively without capturing, sacrificing, or even observing fish, suggests a path toward real-time environmental health monitoring that complements traditional ecological survey methods.

Notably, this integrated readout addresses questions that no single molecular assay can answer alone. Only when interpreted together through the Freshness Gate framework does a fuller picture emerge: which organisms are present, whether they are contributing fresh biological input, how old they are, whether they are healthy, and whether the molecular evidence is temporally reliable. This convergence of independent molecular axes from a single non-invasive sample represents a shift from eDNA as species detection toward environmental nucleic acids as ecosystem diagnostics.

An important distinction underlies these environmental RNA results. The term’eRNA’ is used broadly. Unlike environmental DNA, environmental RNA comprises multiple molecule types with fundamentally different properties. Most eRNA studies since the first applications in aquatic macroorganism systems (Pochon et al., 2017) have detected ribosomal RNA (rRNA), which is highly abundant, structurally stable, and persists longer in the environment than previously assumed (Miyata et al., 2021). Environmental messenger RNA (emRNA), by contrast, is far less abundant and theoretically more transient: it lacks both the extensive secondary structure and ribonucleoprotein association that stabilise rRNA, and the double-helical structure that stabilises DNA (Brandão-Dias et al., 2025), though its detectability depends not only on molecular half-life but also on detection level and continuous production by living organisms (Aminaka et al., 2025). Nearly a decade after the first aquatic eRNA studies, progress remains largely confined to controlled mesocosm experiments demonstrating that emRNA can capture physiological states such as spawning activity, stress responses in tank settings, or life stage (Tsuri et al., 2021; Aminaka et al., 2025; Hechler et al., 2025). While these proof-of-concept studies are necessary, a recent synthesis of the field noted that none of the major eRNA studies specifically investigated eRNA in natural systems, highlighting the persistent gap between theoretical promise and field application (Pochon et al., 2025).

Our detection of spawning-specific mRNA transcripts from environmental water, which was enabled by SISPA amplification of all RNA molecules combined with nanopore sequencing, addresses this gap directly. While specific mRNA has been detected with targeted PCR assays from wild populations, including amphibian life stage determination (Parsley and Goldberg, 2024), our study represents the first recovery of multi-gene functional eRNA profiles (metatranscriptomics) from wild macroorganisms. Previous studies had evaluated ribosomal RNA from multiple organisms before (Littlefair et al., 2022), but the distinction between mRNA and rRNA matters in this context because rRNA presence confirms biological material exists in a sample, while mRNA, with its shorter environmental persistence, provides a more temporally resolved indicator of recent biological input. Yet the power of these molecular signals depends entirely on when they are measured relative to the biological events that generated them.

A single environmental sample, read with shotgun sequencing, is not one measurement but several. From the same library we recover environmental mRNA indicating fresh biological input across multiple functional categories (Table 1), CpG methylation reporting the demographic composition of the DNA pool, pathogen profiles reporting community health, and a housekeeping transcript baseline (histone mRNA) serving as a proxy for total salmon mRNA input. Figure 3 demonstrates this: the same nine weeks of water, viewed through each lens in turn, resolve a coherent picture of the spawning run that no single amplicon assay could provide. Shotgun environmental sequencing recovers functional transcripts tied to ecological events we can independently anchor, and it does so from one non-invasive sample.

This capability comes with a boundary on interpretation that we state plainly: detection is not expression. Environmental water is not a tissue sample, and it is not the sample type built for quantifying gene regulation; because our SISPA workflow amplifies all RNA non-specifically, the transcripts we recover report the presence and relative input of functional molecules, not differential expression among them. When a category such as death reads as zero early in the run, that reflects the absence of biological input (no carcasses yet), not a down-regulated gene. In the same way, methylation reports a state, not an age: we can read the methylation composition of the DNA pool at any time, but it maps to biological age only while fresh DNA dominates, that is, only inside the Freshness Gate. Outside the Gate, degradation decouples methylation from the current biological state, and an age inference would reflect the decay history of the sample rather than the demography of the organisms that shed it.

### 4.2 The Freshness Gate: Temporal boundaries, mechanism, and cross-species validation

Our central finding that environmental methylation reliability exists on a continuum governed by fresh versus degraded DNA concentrations, establishes fundamental constraints on environmental omics inference, extending recent theoretical frameworks for epi-eDNA (Balard et al., 2024). The’Freshness Gate’ is not a fixed temporal window but an operational threshold where biological signal quality depends on sampling timing relative to biological input dynamics. During Coho peak spawning (Weeks 1-3), fresh adult DNA overwhelms background degradation, yielding adult-like methylation indices with narrow confidence intervals. By contrast, later weeks (5, 8) show continued biological input (elevated environmental RNA and qPCR), yet methylation drifts juvenile-ward as accumulated degraded DNA from earlier spawning waves dilutes fresh signals.

While demonstrated here in a freshwater salmonid system with predictable phenology, the Freshness Gate framework should apply to any system where organisms shed nucleic acids in temporally structured pulses, such as marine spawning aggregations, amphibian breeding choruses, migratory stopovers, or coral bleaching events, though threshold parameters will require system-specific calibration.

This finding reconciles an apparent paradox: why do environmental RNA and methylation signals sometimes align (Weeks 1-3) yet other times diverge (Weeks 5, 8)? The answer lies in recognizing that environmental RNA indicates recent biological input (short-lived transcripts with hours-to-days persistence; Yates et al., 2021; Pochon et al., 2017), while methylation reflects cumulative DNA mixtures integrating inputs over weeks (Brandão-Dias et al., 2025; Harrison et al., 2019). Environmental DNA methylation is thus cumulative: early-season samples capture predominantly fresh spawner DNA with adult methylation signatures, whereas later samples accumulate degraded DNA in which methylation marks are progressively lost or rendered undetectable during environmental decomposition.

Two supporting analyses, which we report in the Supplementary Material, are consistent with this reading. A lead-lag analysis shows histone mRNA signals preceding Age Index changes by at least one week (Supplementary Figure S10), consistent with transcript input (hours to days; Cristescu, 2019) arriving before cumulative shifts in DNA composition (days to weeks; Barnes and Turner, 2016). The correlation is significant for 5mC (r = 0.72, p = 0.043) and directionally consistent for 5mCG (r = 0.65, p = 0.082); because sampling was weekly and n = 9, we treat this one-week offset as the minimum detectable lag at our sampling resolution and a supporting observation, not as independent mechanistic proof.

Separately, PERMANOVA variance partitioning indicates that biological freshness (Age Index) accounts for more methylation structure than technical factors (Fragment Length, Read Depth), with directionally consistent partial R^2^ across 5mC and 5mCG, though statistical power is limited (n = 9). Together these are consistent with environmental methylation tracking biological state rather than sequencing artifacts. Resolving the precise timescale of mRNA-to-methylation coupling will require sub-weekly (e.g., daily) sampling during peak spawning activity.

Fragment-length analyses (Supplementary Figure S2) show that the methylation-based age signal is consistent across fragment size fractions above 1 kb within a given week. Across weeks, samples from the degraded phase show lower methylation across these fractions, while fresh-phase samples retain adult-like methylation, indicating that temporal changes in the DNA pool, not fragment length per se, drive the Age Index trajectory. Because fragment length in environmental samples reflects multiple processes (endonuclease activity, mechanical shearing, hydrolysis), it is not a direct proxy for DNA age, and we cannot resolve the temporal ordering of methylation loss relative to fragmentation from these data, though spontaneous deamination of 5-methylcytosine (Lindahl, 1993; Shen et al., 1994) remains a plausible contributing mechanism. Controlled post-mortem studies have shown that DNA degradation increases methylation measurement variance beyond a quality threshold, even when mean methylation remains stable (Rhein et al., 2015); environmental samples face the additional challenge that fresh and degraded DNA pools mix continuously, shifting not only measurement precision but the compositional baseline itself. Characterizing these degradation kinetics at sub-weekly resolution, for example through daily sampling or controlled mesocosm time-series, is an important next step for designing preservation protocols and sampling strategies that maintain interpretability of environmental omics data.

Chinook and Coho salmon showed reproducible Freshness Gate dynamics but with critical differences driven by sampling timing relative to species phenology, consistent with known differences in run timing between these species in Puget Sound tributaries (Quinn, 2018). Chinook exhibited greater methylation heterogeneity because our sampling began after their spawning peak, capturing predominantly late-run and post-spawning degraded signals. Critically, this result independently validates the Freshness Gate model: methylation inference was less informative for Chinook precisely where theory predicts it should, during post-peak sampling when degraded DNA dominates fresh input. The Chinook data thus function as a confirmatory negative control, demonstrating that the fresh:degraded threshold governs demographic inference regardless of species identity. Species-specific molecular half-lives likely govern these dynamics, shaped by physiological traits including body size, lipid content, gamete production, and spawning habitat water chemistry (Jo et al., 2019; Pilliod et al., 2014; Lance et al., 2017). Developing quantitative models linking species traits to environmental DNA and RNA half-lives represents an important next step for predictive environmental omics.

### 4.3 Coupled ecosystem dynamics: Pathogen succession tracks host biology

Pathogen dynamics add ecological context to host molecular signals and complete the multi-omic view of ecosystem state. *Aeromonas* proliferation follows spawning peaks by approximately one week, consistent with opportunistic bacterial growth on spawning-stressed tissue and gamete deposition (Cipriano and Bullock, 2001; Bass et al., 2015). *Saprolegnia* colonization lags by 2-3 weeks, consistent with opportunistic colonization of senescent and necrotic tissue that proliferates during carcass decomposition (Van West, 2006).

Not all classifications survived validation. Kraken2 classified reads tentatively as *Piscine orthoreovirus* (PRV) during mid-season (Weeks 4-6), but alignment to PRV-1 genome segments produced no genome-wide coverage (Supplementary Figure S12), illustrating a key limitation of k-mer-based detection: short sequence matches can produce taxonomic assignments that do not survive alignment. We retain PRV in Figure 3D to show all Kraken2 classifications transparently, while cautioning that its presence remains unconfirmed. That a putative RNA virus was flagged alongside confirmed bacterial and oomycete detections illustrate the breadth of signals recoverable by shotgun meta-transcriptomics, though alignment-based validation remains essential. The high relative abundance of salmon at our sampling location likely facilitated pathogen signal recovery; the approach will be more challenging in more biodiverse or larger systems where host-derived reads are diluted.

Two confirmed microbial classes—bacterial opportunists exploiting stressed tissue and oomycete saprotrophs colonizing necrotic substrates—represent an ordered microbial succession recoverable from environmental nucleic acids. This coordinated temporal cascade, starting from mRNA detection through methylation shifts to pathogen succession, reveals an ecological sequence spanning host physiology to microbial community dynamics on a single molecular platform. Recovering this full ecological cascade without taxon-specific primers or separate assays demonstrates the feasibility of environmental multi-omics for field deployment.

Such integrative capacity suggests opportunities for real-time environmental health monitoring that bridges traditional ecological survey disciplines. Portable nanopore sequencers could be deployed in the field for on-site analysis during critical management periods—salmon runs, harmful algal blooms, disease outbreaks, invasive species incursions (Egeter et al., 2022; Ip et al., 2025b; Maestri et al., 2019). Adaptive sampling allows targeted enrichment of management-relevant taxa while preserving community context, and modified basecalling detects CpG methylation without additional wet-lab steps beyond standard library preparation (Martin et al., 2022; Ni et al., 2024).

However, responsible deployment of environmental omics for management requires conservative interpretation guided by Freshness Gate principles. Methylation-based demographic inference should be restricted to demonstrated fresh-input periods. Convergent environmental mRNA detection and qPCR elevation provide a practical screen for such periods, but as our Week 8 result shows, they do not guarantee reliable inference once degraded DNA accumulates; bootstrap confidence intervals on the Age Index itself should be inspected before demographic conclusions are drawn. Triggered sampling designs that involve deploying intensive omics only after initial detection of biological input or disease signals could optimize cost-effectiveness while targeting Gate Open conditions. Rather than conducting expensive multi-omic sequencing throughout an entire season, managers could use inexpensive qPCR screening to identify Gate Open periods and deploy intensive omics sampling only during those windows, reducing costs while improving the odds of usable data. Future implementations could further streamline this workflow by deriving salmon DNA abundance directly from the shotgun sequencing reads, eliminating the need for a separate qPCR assay while providing an internal concordance measure from the same library.

Integration with remote sensing, hydrological modeling, and population genomics could provide multi-scale context for interpreting molecular signals in complex natural systems.

### 4.4 Limitations and future directions

Two limitations are fundamental to how these data should be read. First, environmental RNA detection is not gene expression. Because environmental water is not a tissue sample and our amplification is non-specific, we can show that a functional transcript is present and track its relative input over time, but we cannot infer that a gene is up-or down-regulated, nor compare expression levels across categories as differential expression.

Second, environmental methylation measures the state of the DNA pool, and that state supports demographic inference only within the Freshness Gate, when fresh DNA dominates. Outside that window, demethylation and fragmentation decouple the signal from current demography, so any age estimate reflects the degradation history of the sample rather than the biology of the organisms that shed it. These are not shortcomings of the sequencing but properties of the sample medium, and recognizing them is what makes the inferences we do report trustworthy.

Our study has several important limitations that constrain generalization and suggest priorities for future work. Weekly sampling resolution limits our ability to resolve sub-week dynamics. The true coupling between environmental RNA and methylation may be tighter than our one-week lag estimate, and daily sampling during peak spawning would better characterize molecular response kinetics. Our sample size (n = 9 weeks) limits statistical power for multivariate analyses; PERMANOVA tests showed consistent effect directions but did not reach significance, and replication across multiple watersheds and years is needed to establish robust effect sizes. qPCR measurements were available only from Week 4 onward, so Gate Open classification for the earliest weeks rests on environmental RNA alone.

Generalization to other taxa, ecosystems, and environmental media (sediment, seawater, air, soil) requires systematic testing of Freshness Gate parameters beyond the single watershed and two closely related salmonid species examined here.

Our Age Index calibration relies on a single reference per life stage and species: adult muscle tissue from one wild-caught individual per species, and juvenile eDNA from rearing-tank water. While the large number of age-informative CpG sites (364-1,100 depending on species and modification type) provides dimensionality comparable to multi-individual SNP panels used in population assignment, biological replication of calibration tissues from multiple individuals across different populations would strengthen confidence in the adult-juvenile methylation axis. The current calibration establishes a relative ordering (more adult-like vs. more juvenile-like) rather than an absolute demographic assignment, and future studies should expand calibration panels to include intermediate life stages (eggs, alevins, fry, smolts) for finer-scale demographic resolution. We did not orthogonally validate methylation calls with bisulfite sequencing or other gold-standard methods; while tissue calibration samples showed expected adult-juvenile separation and environmental patterns were biologically consistent, independent validation would strengthen confidence in absolute methylation levels. Future studies should also explore whether RNA stabilization protocols (spike-ins, immediate preservation and extraction) can extend environmental RNA interpretability beyond the short windows we observed, and whether controlled mesocosm studies manipulating temperature, flow, and pH could quantify the effects of environmental covariates on molecular half-lives.

During the study period, a sewer leak from private property was discovered flowing into Issaquah Creek upstream of the sampling site on September 25, 2024 (City of Issaquah, 2024), one day before our Week 5 sampling event. The leak was repaired within days and the creek advisory lifted by October 3. While elevated human-associated microbial taxa were detected in post-event weeks, disentangling sewage effects from seasonal increases in rainfall and organic loading from carcass decomposition is beyond the scope of this study.

### 4.5 Molecular clocks in the environment: A new framework for ecological inference

By establishing that different environmental nucleic acid molecules have characteristic degradation timescales—from hours for transcripts, through days for methylation reliability, to days-weeks for bulk DNA presence—our work reveals a biological clock embedded in environmental samples. Though not yet calibrated to the quantitative precision of radiometric dating or dendrochronology, this temporal structure provides an analogous conceptual framework for retrospective inference and prospective monitoring design. Just as archaeologists use isotope ratios to date artifacts, ecologists may eventually use environmental RNA:DNA ratios, methylation state distributions, and fragment length patterns to infer when biological inputs occurred and whether environmental samples still faithfully represent current ecosystem state.

Environmental DNA thus moves from a static snapshot technology to a dynamic record of ecosystem change. Rather than asking’what is present?’, we can now ask’what is happening?’,’how old is this biological signal?’, and’when should we trust molecular inferences?’. By defining the Freshness Gate and mapping its empirical boundaries, we establish practical and theoretical boundaries for environmental omics monitoring.

Environmental nucleic acids are not permanent molecular fossils but living biological information that expires on timescales from hours to weeks. Recognizing these expiration dates turns a limitation into useful information. With the understanding of when molecular signals expire, we can design sampling strategies that capture reliable biological information, and read not just what lives in an ecosystem but when its molecular signals can be trusted and what they reveal.

## Acknowledgments

We thank the Issaquah Salmon Hatchery staff for site access and hatchery escapement data. We are grateful to Ian Young of the Seattle Aquarium for facilitating collection and filtration of juvenile salmon aquarium holding tank water. We thank Prof. Lauren Sassoubre for helpful feedback on environmental RNA interpretation. We also acknowledge OceanKind [Grant No. GR042190] and the David and Lucile Packard Foundation [Grant No. GR016745] for funding support and the Center for Environmental Genomics for access to HPC resources.

## Author Contributions

Y.C.A.I. designed the study with input from E.A.A. Y.C.A.I., G.G., and P.F.P.B.D. conducted field sampling and laboratory work. Y.C.A.I. performed bioinformatic analyses.

Y.C.A.I. and G.G. conducted qPCR assays. Y.C.A.I. and R.P.K. wrote the manuscript.

E.A.A. contributed to manuscript revision. All authors reviewed and approved the final version.

## Data Availability

Raw sequencing data (Bam and FASTQ files) have been deposited in the NCBI Sequence Read Archive under BioProject accession PRJNA[XXXXXX]. Processed datasets including methylation frequency matrices, transcript detection tables, and qPCR data, together with analysis code for methylation calling, transcriptome analysis, and visualization, are available from the corresponding author on request during peer review. Both will be released publicly at https://github.com/piedna/Wild-Omics under MIT license, and archived at Zenodo, upon publication. Reference methylation profiles for age-informative CpG sites are available from the corresponding author on the same basis. Hatchery escapement data are publicly available from Washington Department of Fish and Wildlife (https://wdfw.wa.gov/fishing/management/hatcheries/escapement).

## Ethics Statement

Water sampling was conducted by Washington Department of Fish and Wildlife, Friends of Issaquah at Issaquah Hatchery. No IACUC approval is required as no live animals were handled or sacrificed for this study. All biological material was recovered from environmental water samples. Adult tissue samples for methylation calibration were obtained from commercially available wild-caught salmon purchased at retail. Juvenile calibration samples consisted of tank water from the Seattle Aquarium (no animal handling required).

## Supplementary Materials

Supplementary Materials S1 and S2 are available from the corresponding author on request.

**Supplementary Material S1.** Modified phenol-chloroform-isoamyl alcohol (PCI) DNA extraction protocol for native nanopore methylation sequencing from DNA/RNA Shield-preserved environmental water samples.

**Supplementary Material S2.** Complete list of transcripts detected from environmental RNA in Coho and Chinook salmon. All genes detected at ≥1 read across sampling weeks, organized by functional category with NCBI accessions and references.

---

**Supplementary Figure S1.**
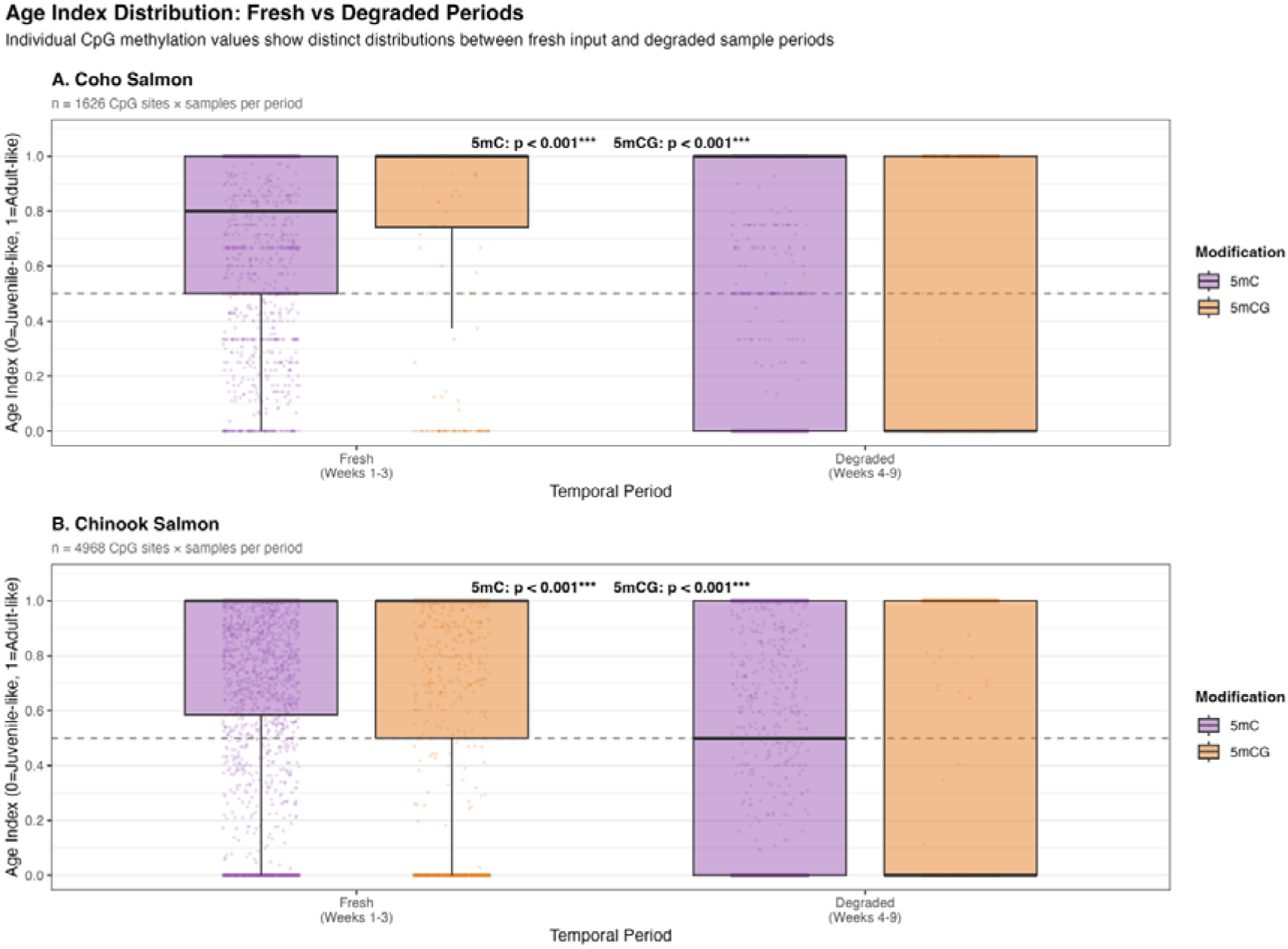
Age Index distribution across fresh and degraded periods. Boxplots of methylation (CpG) Age Index illustrate clear separation between early fresh-input weeks (1-3) and later weeks (4-9) in both Coho and Chinook salmon.

**Supplementary Figure S2.**
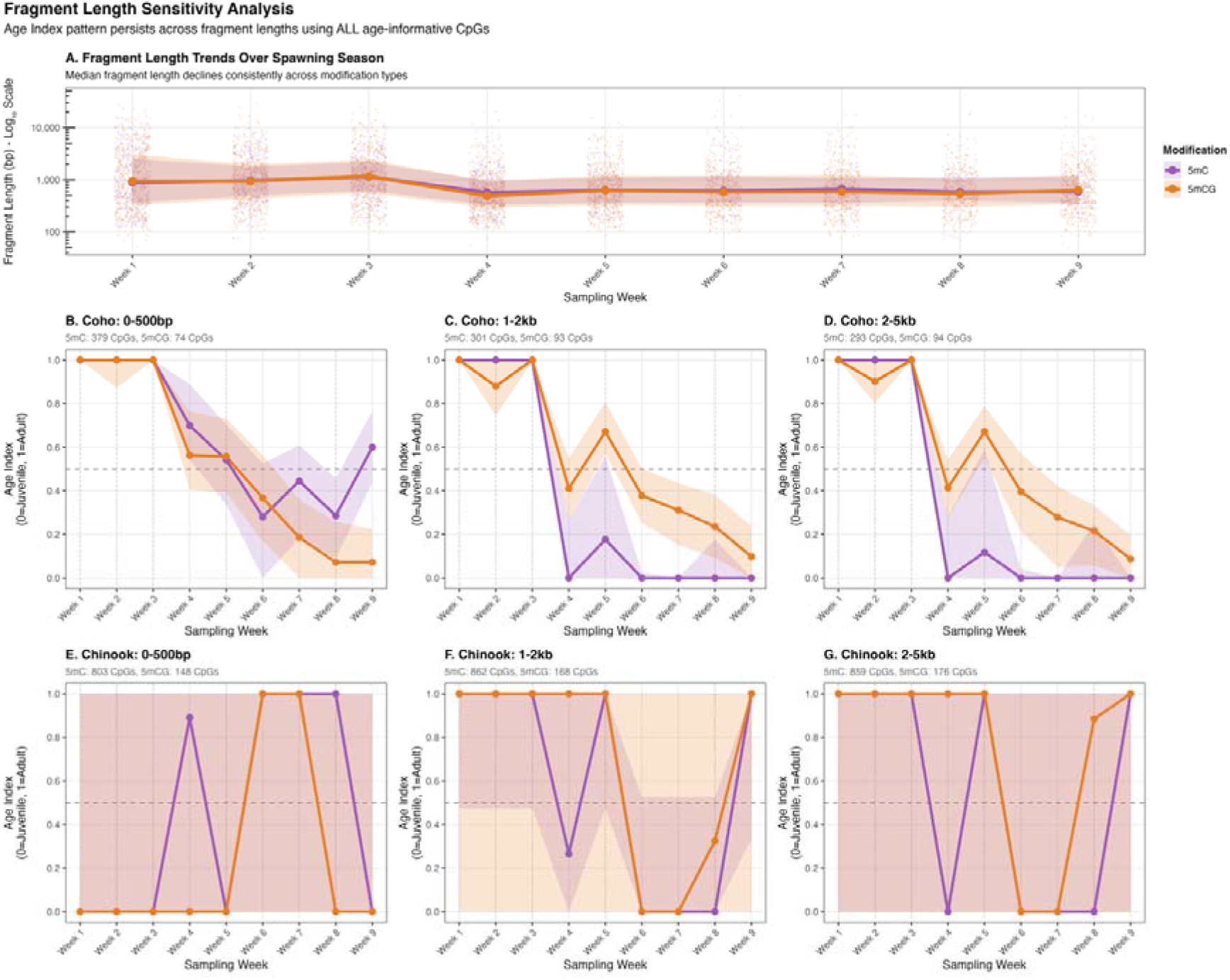
Fragment length sensitivity analysis. Methylation-based Age Index patterns are consistent across fragment length bins above 1 kb in Coho (1-2 kb, 2-5 kb); the 0-500 bp bin and all Chinook bins show substantially greater variability.

**Supplementary Figure S3.**
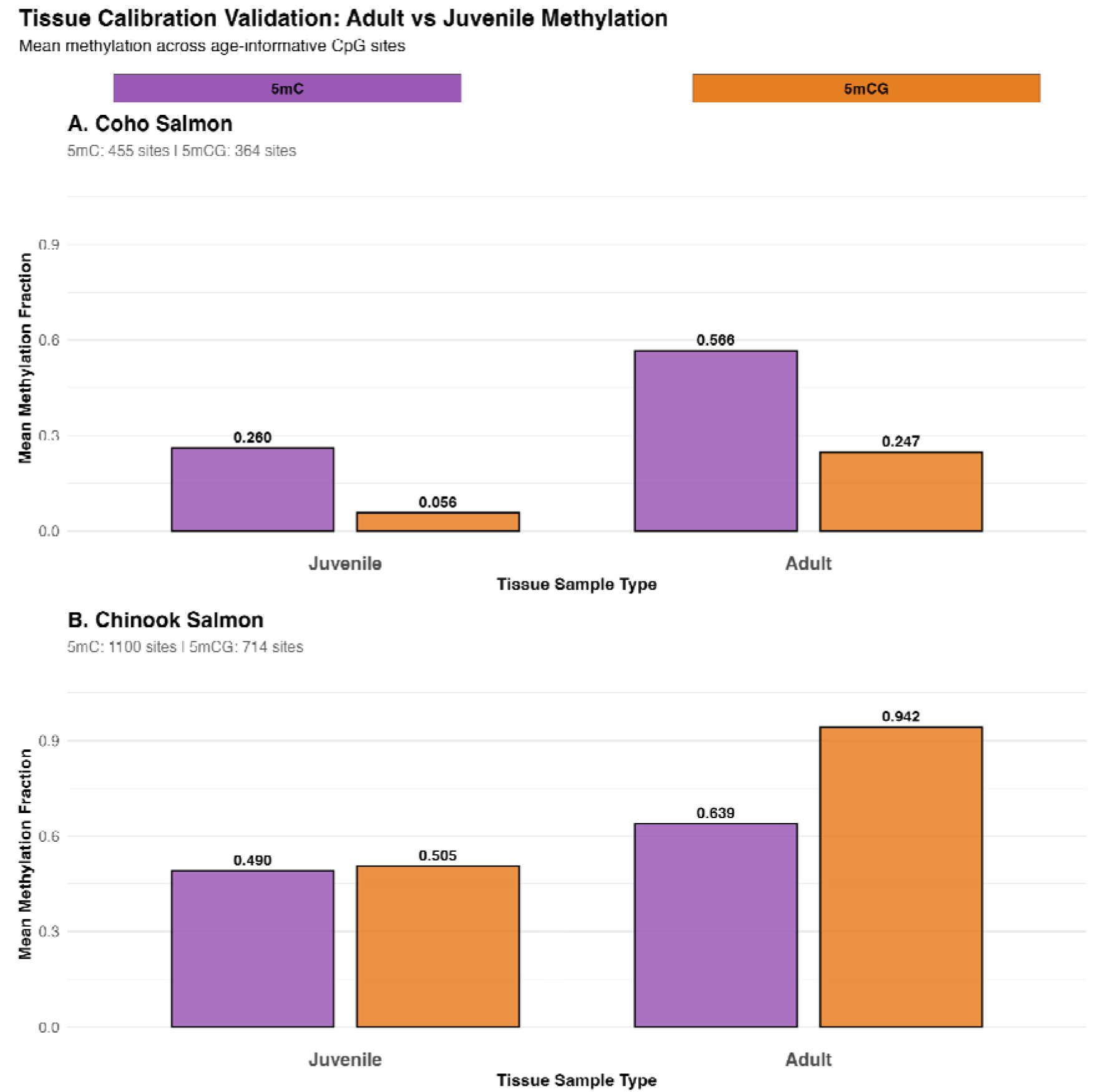
Tissue calibration validation: Adult vs juvenile methylation. Mean methylation fractions across age-informative CpG sites confirm biological separation between juvenile and adult calibration references in both species.

**Supplementary Figure S4.**
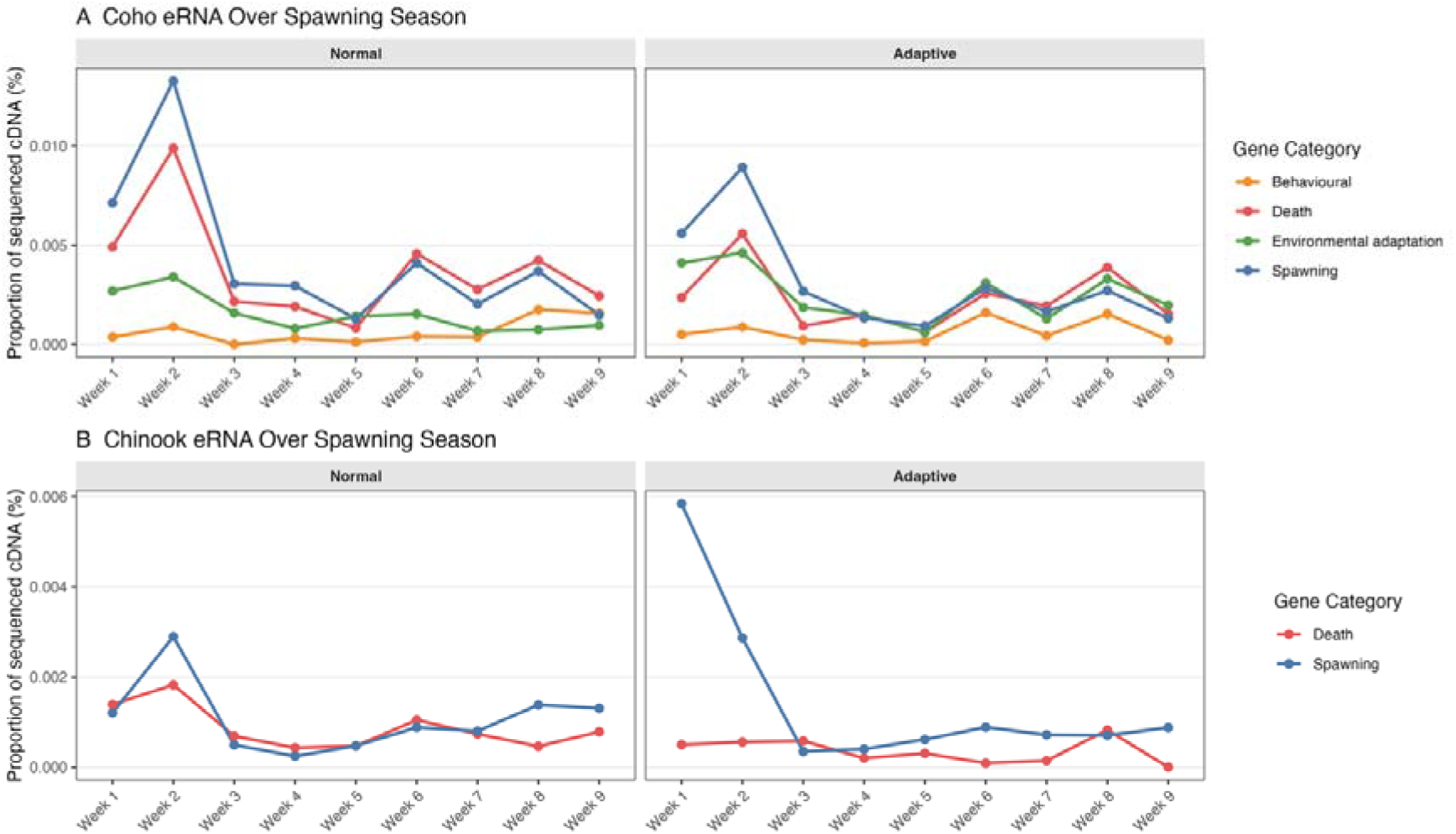
Adaptive sampling improves species assignment. Comparison of environmental RNA transcript detection between normal and adaptive sampling shows improved taxonomic resolution and species-specific signal recovery.

**Supplementary Figure S5.**
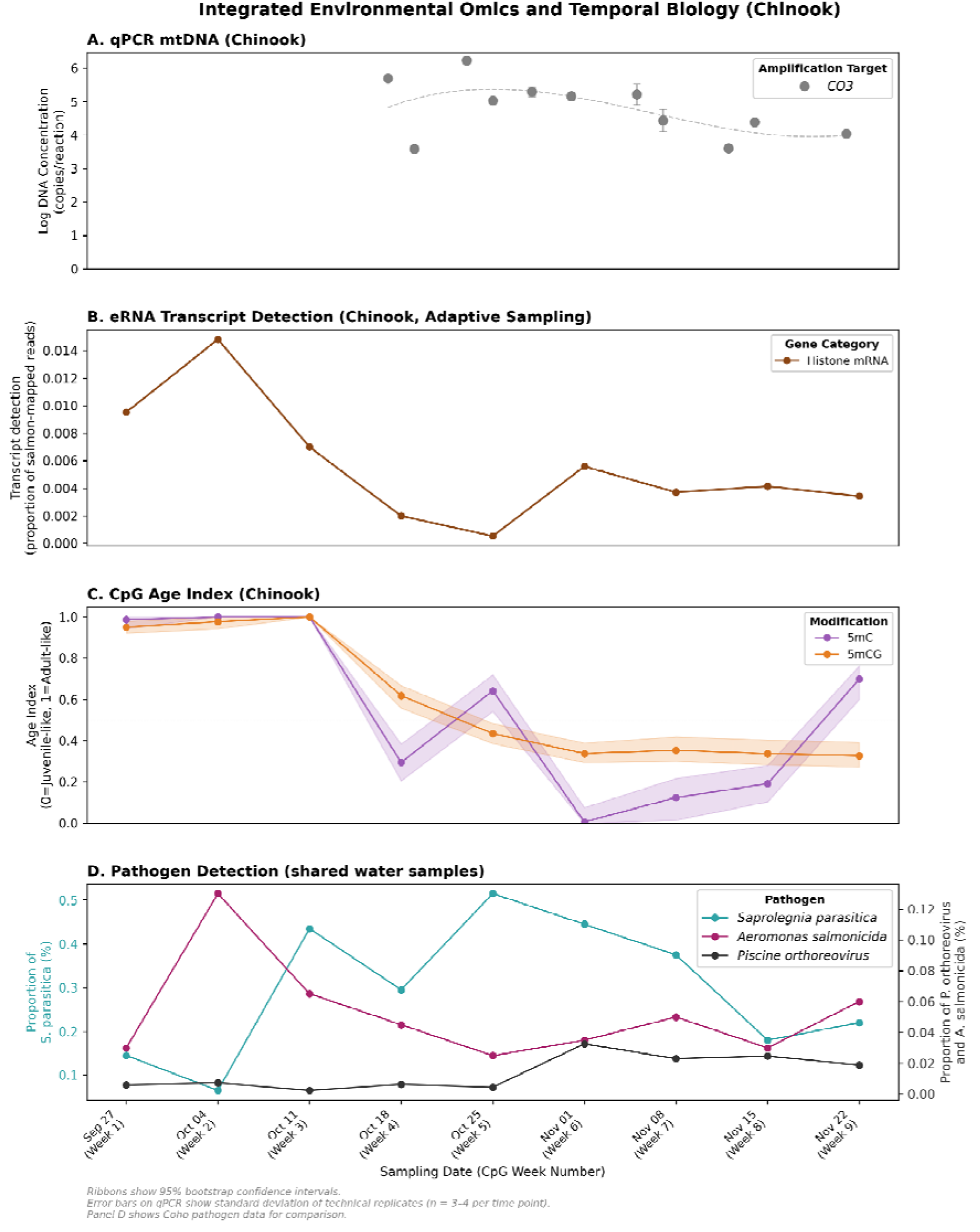
Integrated environmental omics and temporal biology for Chinook salmon. (A) qPCR mtDNA (COX3) concentration from the same water samples; error bars show standard deviation of technical replicates (n = 3-4 per time point). (B) Histone mRNA recovery, expressed as the proportion of salmon-mapped reads, across the sampling period. Chinook eRNA was available for weeks 1 and 5 through 13, producing a temporal gap between 30 August and 27 September. (C) CpG methylation Age Index for 5mC and 5mCG modifications, with 95% bootstrap confidence intervals (ribbons). (D) Pathogen detection from the total metagenomic fraction of the shared water samples (species-independent); data are the same as in Figure 3D because pathogens are detected from untargeted reads, not from species-specific captures.

**Supplementary Figure S6.**
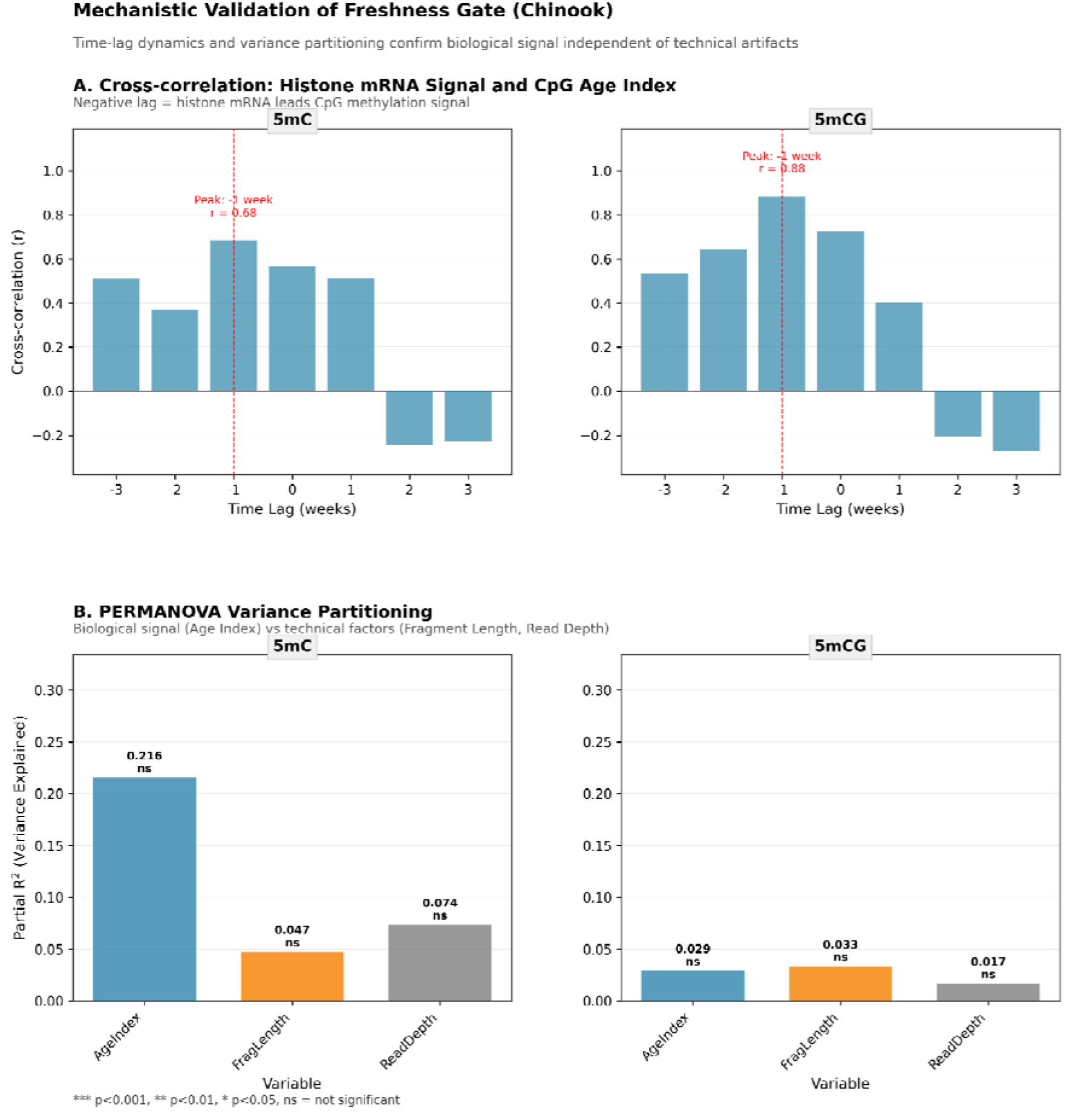
Mechanistic validation of the freshness gate (Chinook). (A) Cross-correlation between histone mRNA signal and CpG Age Index at integer week lags: both 5mC and 5mCG peak at lag =-1 (histone leads by one week; 5mC r = 0.68, 5mCG r = 0.88), consistent with transcript input preceding measurable change in the DNA methylation pool. (B) PERMANOVA variance partitioning of biological versus technical drivers of methylation structure: Age Index explains the most unique variance for 5mC (partial R^2^ = 0.216), above Read Depth (0.074) and Fragment Length (0.047); none reach significance with the available sample size. Results parallel the Coho analysis (Figure 3).

**Supplementary Figure S7.**
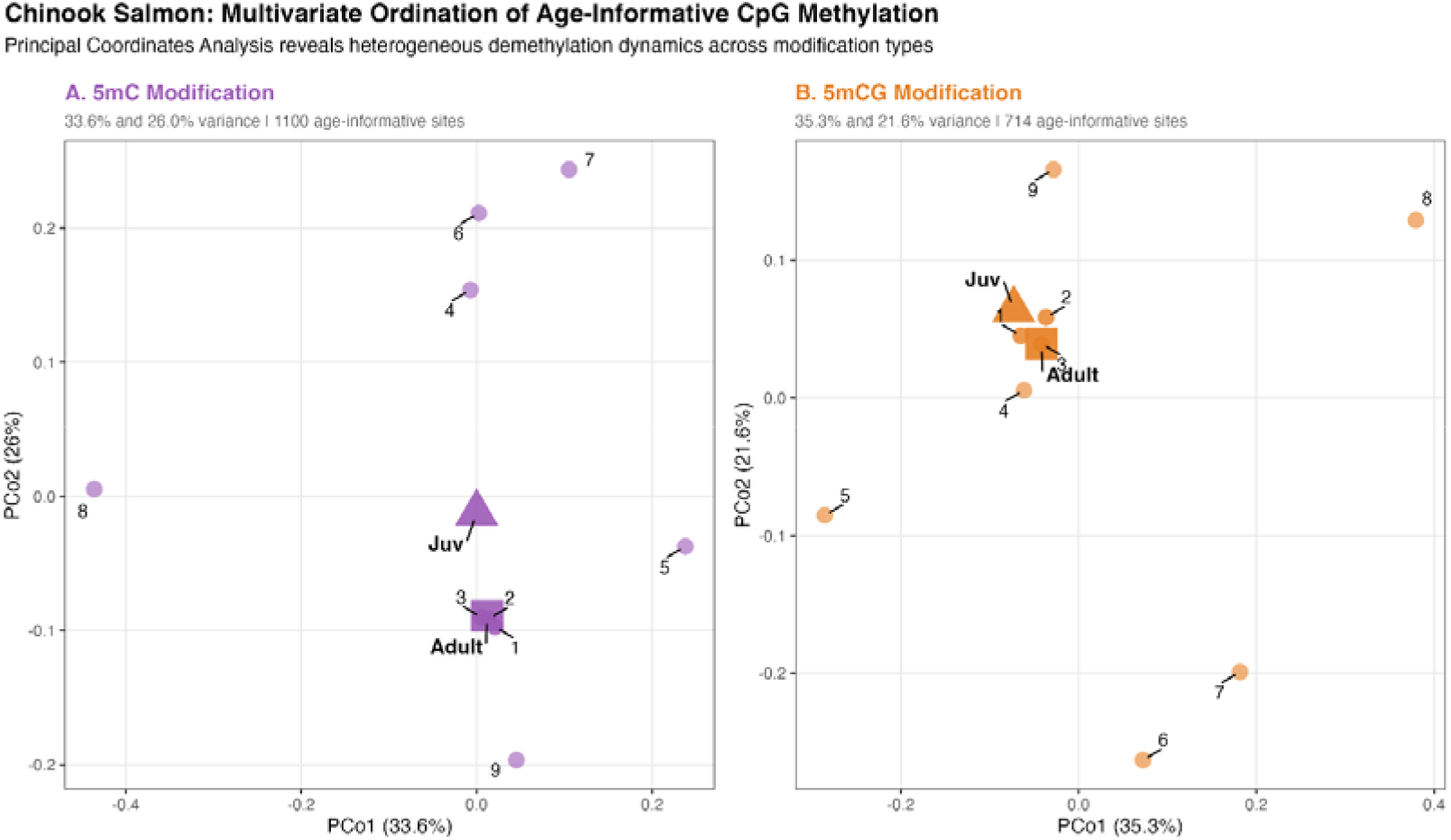
Chinook salmon PCoA of age-informative CpG methylation. Multivariate ordination reveals heterogeneous temporal demethylation dynamics reflecting post-peak sampling design.

**Supplementary Figure S8.**
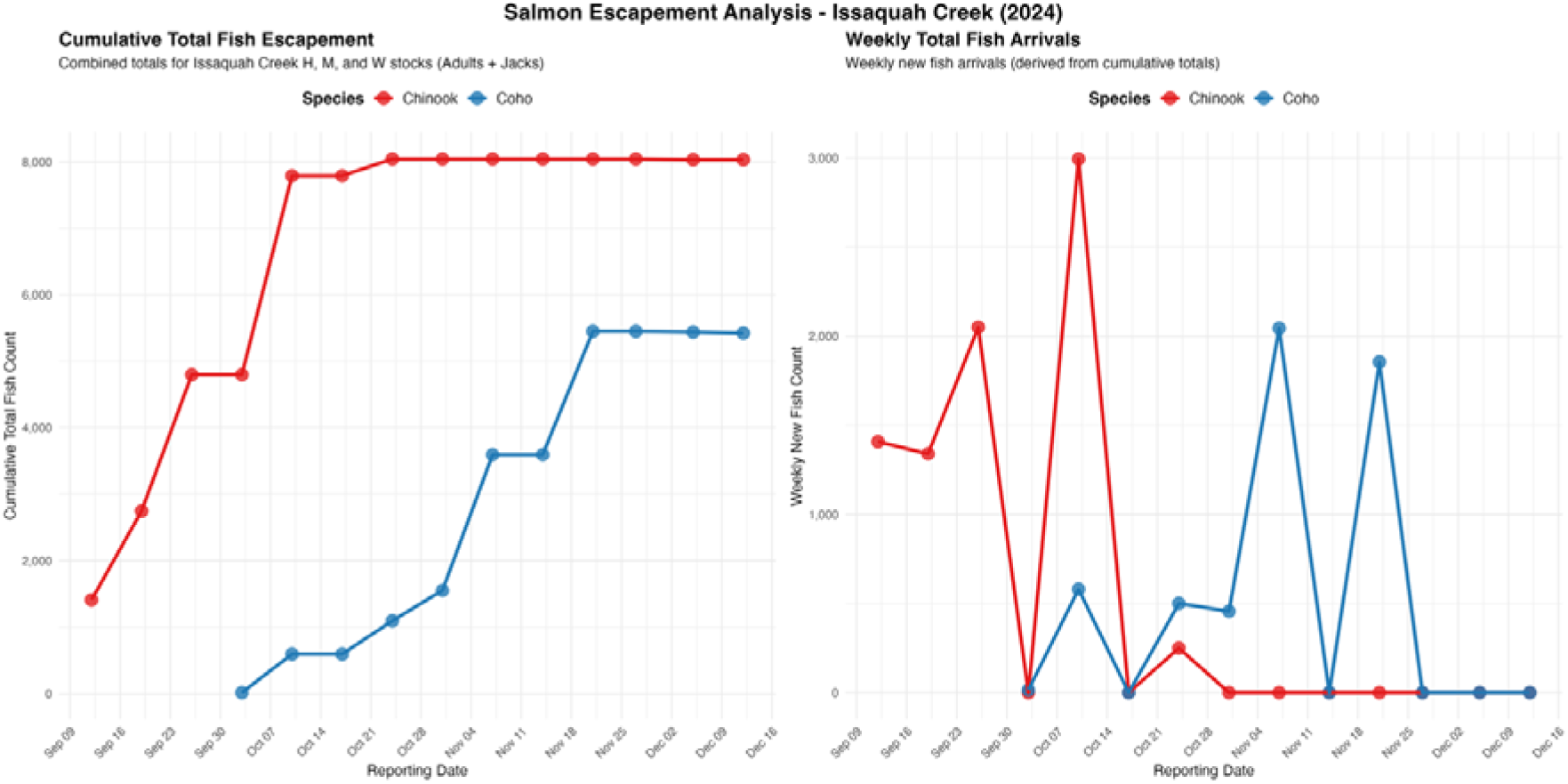
Hatchery escapement reports from Issaquah Creek (2024). Chinook returns declined through October while Coho arrivals extended into November, reflecting distinct run timing between species.

**Supplementary Figure S9.**
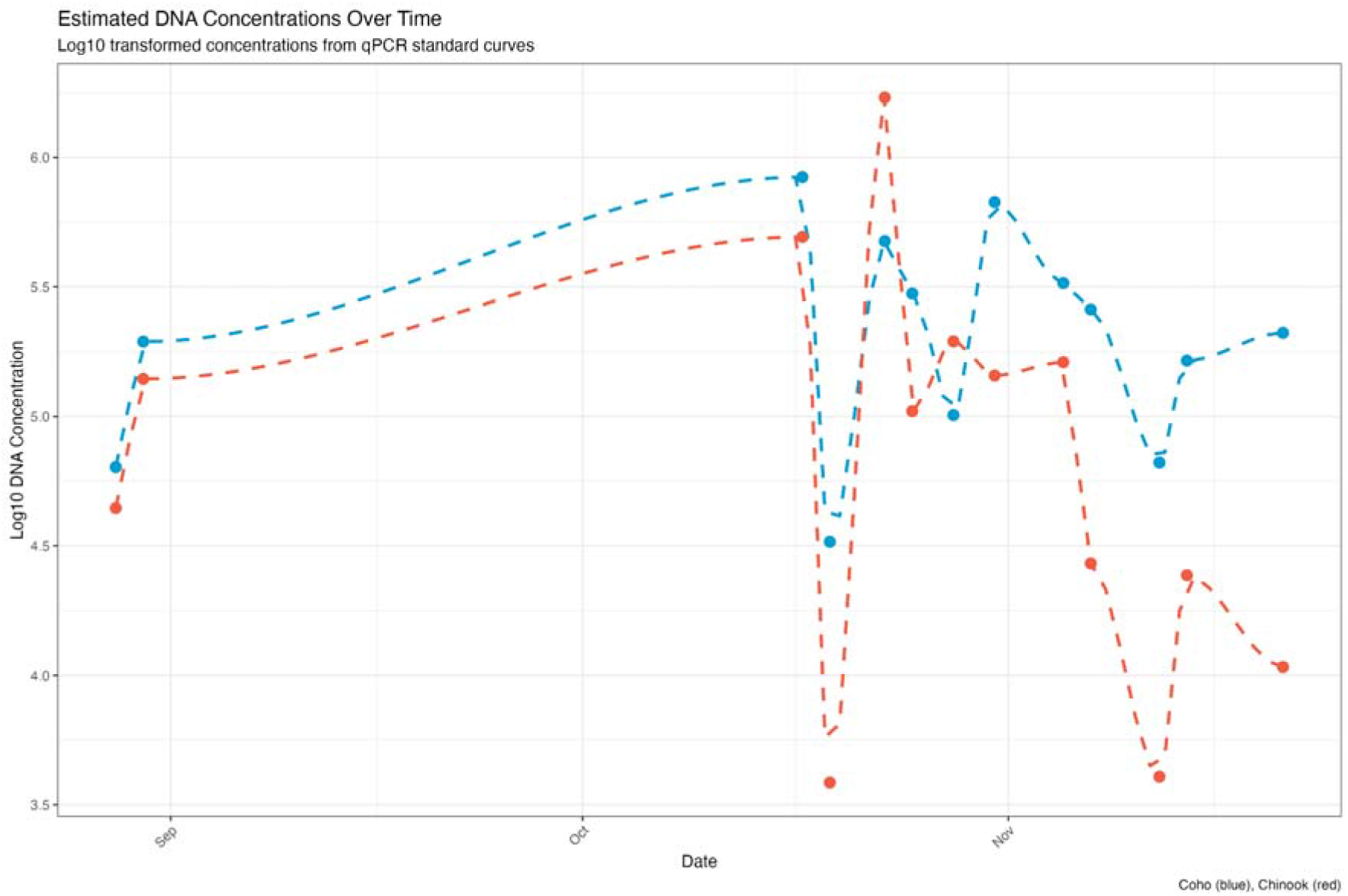
qPCR standard curves and detection probability. Performance characterization of Coho cytochrome b and Chinook COIII/ND3 assays including amplification efficiency, limit of detection, and observation-level uncertainty.

**Supplementary Figure S10.**
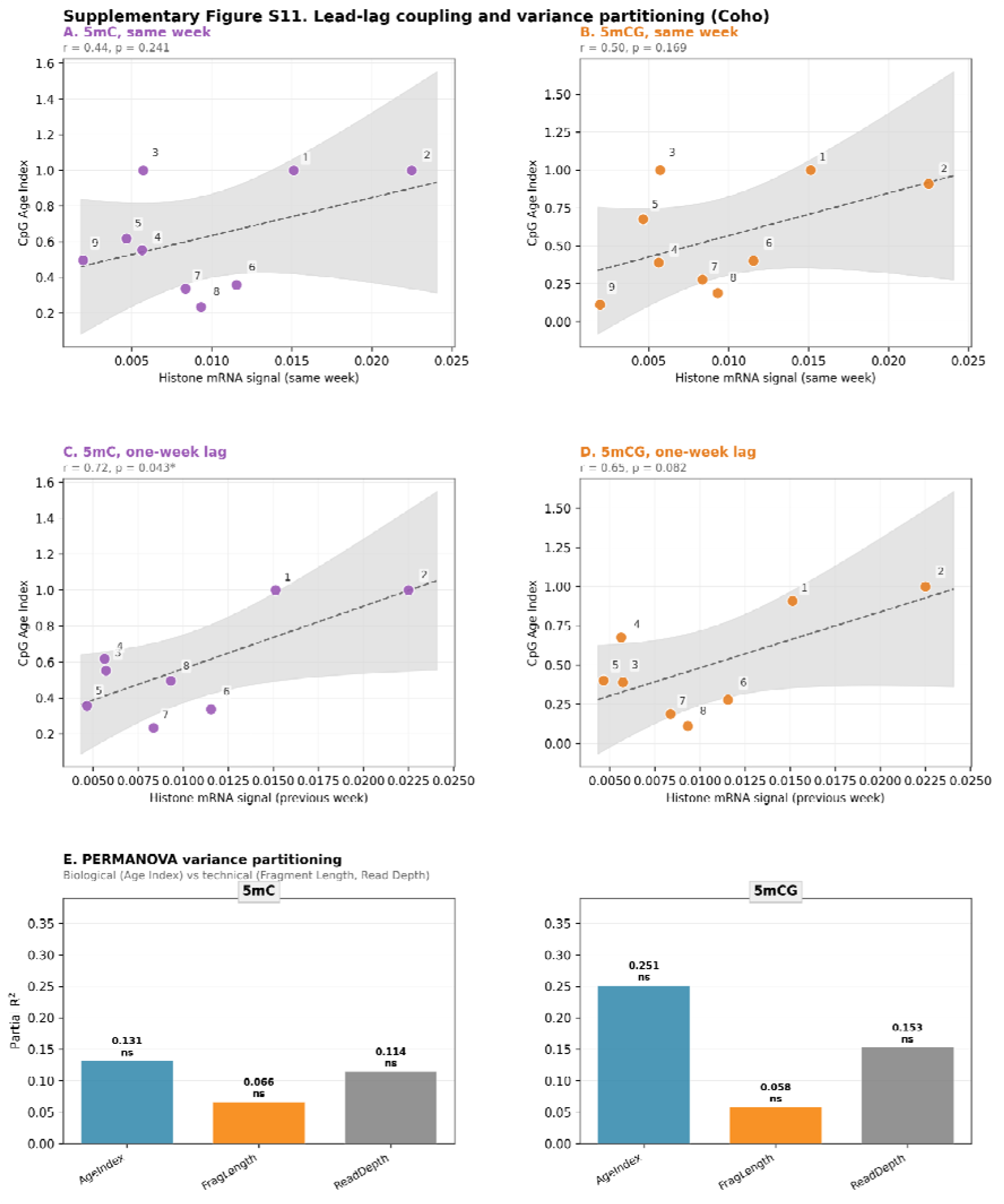
Lead-lag coupling and variance partitioning (Coho). (A, B) Histone mRNA proportion versus CpG Age Index in the same week: positive but not significant (5mC r = 0.44, p = 0.241; 5mCG r = 0.50, p = 0.169). (C, D) With histone mRNA shifted forward one week the relationship strengthens (5mC r = 0.72, p = 0.043; 5mCG r = 0.65, p = 0.082), consistent with transcript input preceding change in the DNA methylation pool; because sampling is weekly this one-week offset is a minimum detectable lag. (E) PERMANOVA of biological versus technical drivers of methylation structure: Age Index explains the most unique variance (partial R^2^: 5mC 0.131, 5mCG 0.251), above Read Depth (0.114; 0.153) and Fragment Length (0.066; 0.058); none reach significance.

**Supplementary Figure S11.**
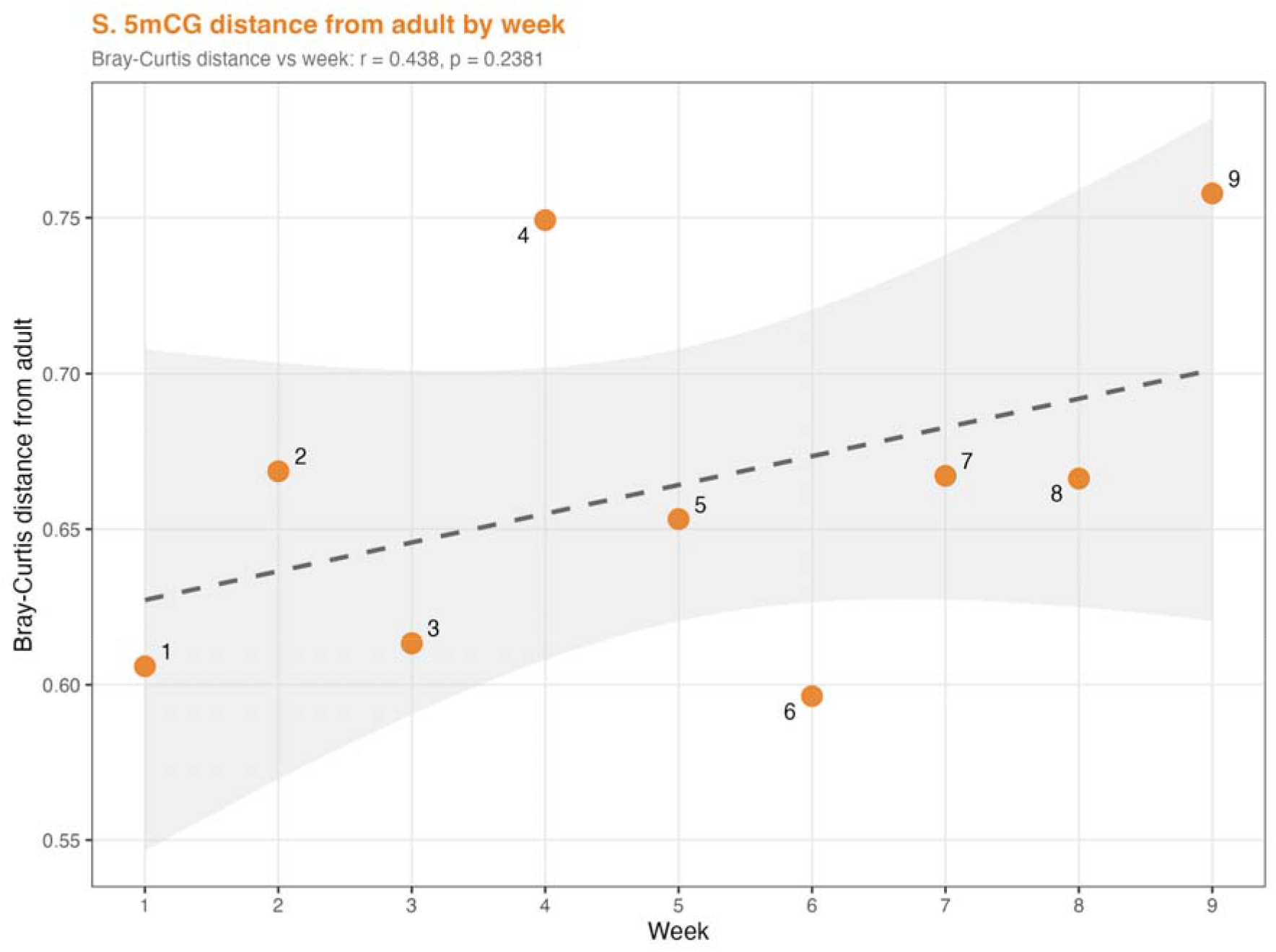
Bray-Curtis distance from the adult calibration sample by week for 5mCG shows dispersion rather than directional drift (r = 0.438, p = 0.238), in contrast to the progressive 5mC trend shown in Figure 4C.

**Supplementary Figure S12.**
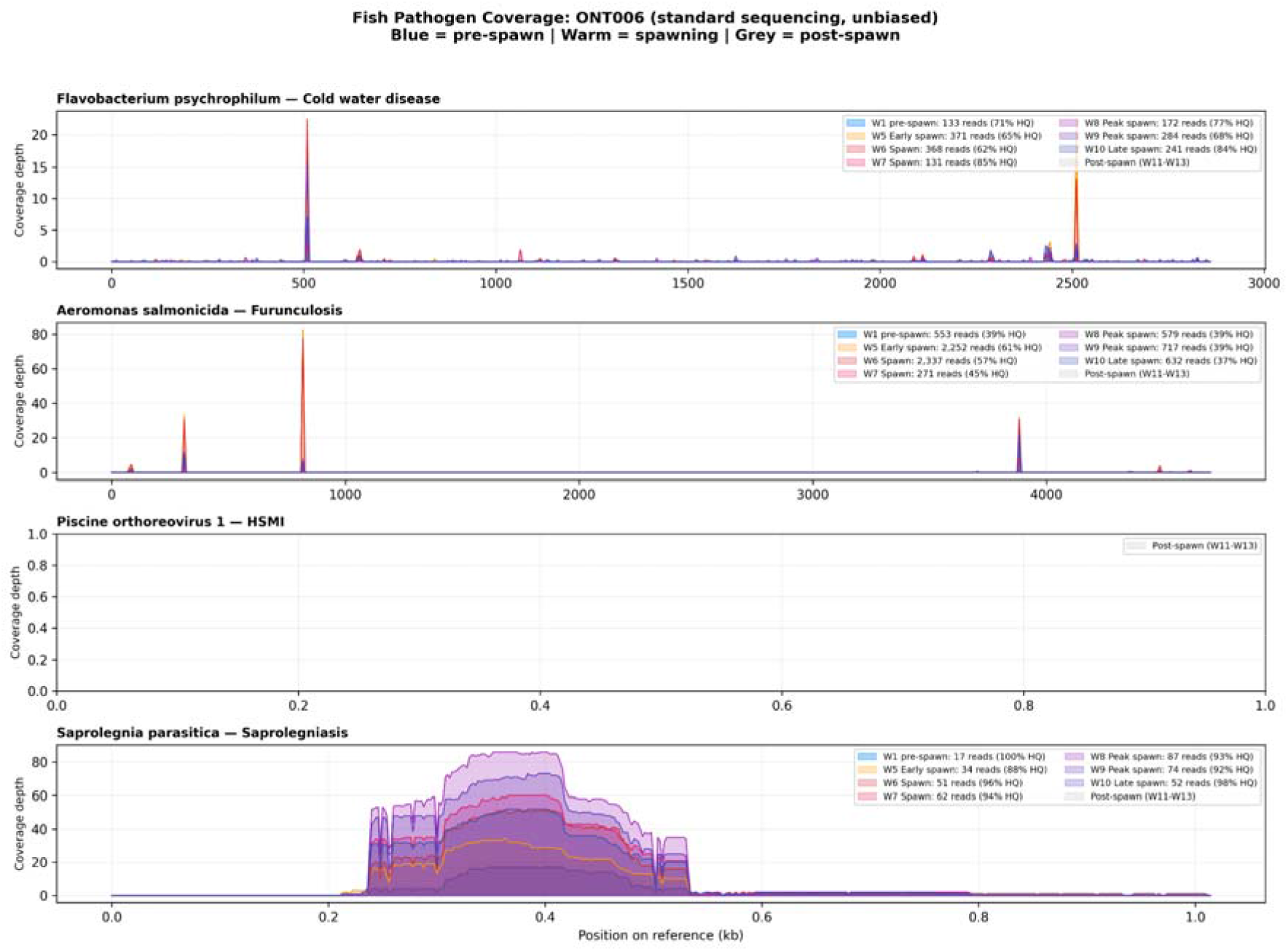
Minimap2 alignment validation of pathogen detection. Genome-wide read coverage for four fish pathogens, with reads aligned to reference genomes (*F. psychrophilum* NC_009613.3, *A. salmonicida* NC_009348.1, PRV-1 genome segments, *S. parasitica* ITS/actin markers). Color gradient from blue (pre-spawn) through warm tones (spawning) to grey (post-spawn) shows temporal patterns consistent with Kraken2-based classifications (Figure 3D). PRV-1 shows no aligned reads despite Kraken2 classification, suggesting that k-mer matches may reflect shared sequence features with related viral taxa rather than true PRV presence.

